# The AMPK-related kinase NUAK1 regulates neuronal morphogenesis through the RNA splicing co-factor SON

**DOI:** 10.1101/2025.10.10.681550

**Authors:** Martijn Kerkhofs, Géraldine Meyer-Dilhet, Hélène Polvèche, Sozerko Yandiev, Jacqueline Tait-Mulder, Sergio Lilla, David Sumpton, Daniel J. Murphy, Eun-Young Erin Ahn, Cyril F. Bourgeois, Evelyne Goillot, Julien Courchet

## Abstract

In recent years, alternative splicing emerged as a major mechanism controlling gene-regulatory networks during brain development, yet how alternative splicing is tuned to the dynamic alterations underlying neuronal maturation remains poorly understood. Here, we identified that NUAK1, an AMPK-related kinase linked to neurodevelopmental disorders, is a key regulator of alternative splicing in developing cortical neurons. Mechanistically, NUAK1 exerts its function through phosphorylation of the splicing co-factor SON, regulating a group of highly conserved splicing events in genes crucial for neurodevelopment. We demonstrate that SON plays an important role in cortical neuron development, which is consistent with the neurodevelopmental phenotypes observed in Zhu-Tokita-Takenouchi-Kim (ZTTK) syndrome, a genetic disorder caused by SON haploinsufficiency. Together, our findings uncover a novel pathway involving NUAK1 and SON, which orchestrate a splicing program required for proper neuronal development.

## INTRODUCTION

Neurodevelopment is a complex process that is strictly regulated in time and space. The sequential steps leading to the building of functional neural circuits are under the control of gene-regulatory networks, which have been extensively characterized over the past two decades (*1*). Transcriptomic and epigenomic approaches at the single cell level revealed the dynamic pattern of gene-expression regulation and its critical importance to govern cell fate and morphogenesis in the normal and pathological course of human brain development (*2–5*). Whereas transcription factors have long been identified as markers of neural cell identity and essential components of the neurodevelopmental program, it is now acknowledged that the subsequent steps in the complex cascade of gene expression also play an oversized role in the fine-tuning of neuronal development.

Over the past decade, alternative RNA splicing has emerged as an important node regulating virtually all aspects in neuronal development (*6–11*). In the nervous system, alternative splicing is cell type-specific (*12–14*) and is tightly temporally controlled (*15*). The importance of alternative splicing is highlighted by the identification of a number of splicing factors among the genes associated with neurodevelopmental disorders (*16–18*). This is best exemplified by the recent discovery of *de novo* mutations in the small nucleolar RNA *RNU4-2*, a key component of the splicing machinery, which play a major role in neurodevelopmental disorders (*19, 20*) In addition, in complex genetic disorders such as autism spectrum disorder (ASD), splicing patterns in the brain are significantly disrupted (*21–23*). While we know that alternative splicing is important for proper neuronal differentiation, the mechanisms ensuring a dynamic regulation of the splicing machinery, and adaptation to the changing cellular environment during the course of brain development remain largely unknown.

In this study, we focus on NUAK1, a serine/threonine kinase belonging to the AMPK-related kinase family (*24*), which was identified as a major regulator of terminal axon branching and branch stabilization, both *in vitro* and *in vivo (25-27)*. NUAK1-deficiency results in impaired social behaviours in heterozygous NUAK1 mice (*26*). The phenotypic effects of NUAK1 involve the disruption of a range of mitochondria-associated biology, from localization to function, stemming from the regulation of the mitochondrial micro-protein BRAWNIN at the RNA level (*27*). Furthermore, a recent study implicated NUAK1 in the regulation of RNA splicing in cancer cells (*28*). This led us to investigate whether NUAK1 would have a wider effect on splicing in developing neurons. Our results reveal that NUAK1 is localized in the nuclear speckles, where it is a crucial regulator of alternative splicing in developing cortical neurons by functionally interacting with SON, a ubiquitously expressed splicing regulator and a structural component of the nuclear speckles, where pre-mRNA splicing takes place (*29*). Genetic mutations in *SON* give rise to the Zhu-Tokita-Takenoushi-Kim syndrome (OMIM 617140), a recently discovered, rare neurodevelopmental condition with an important component of brain malformations, intellectual disability and ASD-like behaviour (*17, 30–32*).

Here, we demonstrate that NUAK1 directly phosphorylates SON and affects the structure of nuclear speckles. Functionally, SON and NUAK1 co-regulate highly conserved splicing events occurring throughout different stages of neuronal maturation. Importantly, the NUAK1/SON shared splicing signature is enriched in a subset of genes that are heavily implicated in neurodevelopmental diseases. Finally, we report that SON-deficiency partially phenocopies the lack of NUAK1 in primary cortical neurons at various stages of development. In short, we propose that the regulation of SON by NUAK1 constitutes a novel mechanism through which alternative splicing is temporally controlled during various stages of neurodevelopment.

## RESULTS

### NUAK1 regulates mRNA splicing in primary cortical neurons

The AMPK-related kinase NUAK1 is an essential regulator of cortical axon development (*25–27*). A previous report described a nuclear localization of NUAK1 in cancer cells, where it colocalizes with components of the spliceosome (*28*). We expressed a Flag-tagged NUAK1 in 293T cells, and performed colocalization assays with SON, confirming its localization in nuclear speckles (*29*) (**Suppl. Fig. 1A**). Nuclear speckles are largely accepted as sites regulating mRNA splicing in eukaryotic cells. In light of NUAK1 localization to nuclear speckles in immature cortical neurons, we sought to explore how it controls a broader mRNA splicing program during cortical development. To do so, we performed an acute lentivirus-mediated knockdown of *Nuak1* in primary culture of mouse cortical neurons and harvested the RNA for deep bulk RNA-sequencing at 5 days *in vitro* (DIV) (**Suppl. Fig. 1B-C**). Analysis of splicing using rMATS (*40*) revealed that *Nuak1* knockdown significantly altered 1018 splicing events spread across 813 different genes (**Fig. 1A**). Most NUAK1-controlled splicing events concerned cassette exons: over half of the significant events affected exon inclusion or skipping (ES; 55.4%), or mutually exclusive exons (MXE; 23.6%) (**Fig. 1B**). Intron retention constituted approximately 10% of altered splicing events (IR; 9.1%). Furthermore, knockdown of *Nuak1* resulted in a shift towards more exon skipping and more intron retention, owing to a splicing bias across ES and IR events (**Fig. 1C-D**; **Suppl. Fig. 1D**). To gauge the significance of NUAK1-regulated splicing events, we determined which part of the mature RNA would be altered: the 5’-untranslated region (UTR), the coding sequence of the protein or the 3’-UTR. This analysis revealed that approximately 70% of the observed splicing events would impact the coding sequence of mRNAs (**Fig. 1E)**. Altogether, these results suggest that NUAK1-mediated splicing events could have a significant impact on the proteome in early developing neurons. To gain insight into the biological processes that are affected by splicing changes upon *Nuak1* knockdown, we performed an enrichment analysis for the mis-spliced genes using gProfiler (*41*) (**Fig. 1F**). Alongside known NUAK1-associated processes like regulation of cell morphogenesis, NUAK1-controlled splicing also affects a number of genes important for synaptic function such as post-synapse organization, vesicle-mediated transport in the synapse and maintenance of synaptic structure (**Fig. 1F**). Surprisingly, compared to its substantive effect on splicing, *Nuak1* knockdown only altered the expression of 142 genes (110 upregulated; 32 downregulated) (**Suppl. Fig. 1E**). Only 6% of the genes in the differential gene expression profile were also affected by NUAK1-associated splicing (**Fig. 1G**). Different gene sets and pathways were enriched in the DGE profile compared to the enrichment analysis for splicing (**Suppl. Fig. 1F**), with a significant enrichment in metabolic processes. We conclude that NUAK1 is an important regulator of splicing in developing cortical neurons, impacting genes broadly regulating neuronal development and function. Moreover, NUAK1 seems to differentially regulate gene expression and splicing.

**Figure 1.**
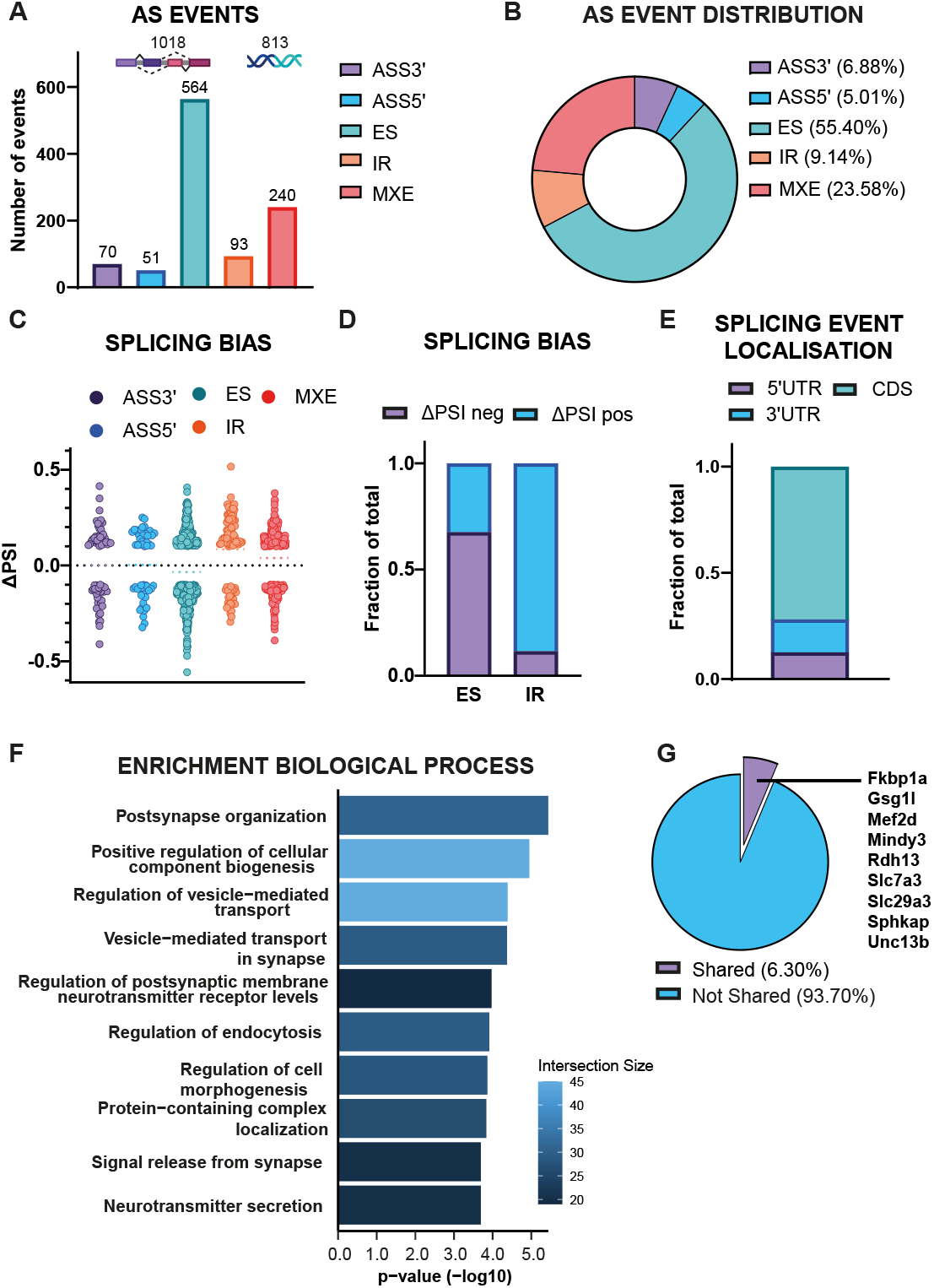
NUAK1 regulates splicing in primary cortical neurons. (A) Number of alternative splicing (AS) events per category compared to the control condition in NUAK1-deficient cortical neurons (5DIV): alternative splice site 3’ (ASS3’), ASS5’, exon skipping or inclusion (ES), intron retention (IR) and mutually exclusive exons (MXE). (B) AS event distribution per category in NUAK1-deficient cortical neurons. (C) Scatter plot showing the distribution of Δ percent spliced in (PSI) per splicing event category in NUAK1-deficient cortical neurons. (D) Splicing bias in splicing events in NUAK1-deficient neurons as measured by the ratio of positive and negative ΔPSI. (E) Splicing event topology in NUAK1-deficient cortical neurons. (F) gProfiler enrichment analysis for biological process on genes that were significantly differentially spliced in NUAK1-deficient cortical neurons. (G) Overlap between the genes found in the AS analysis and the genes found in the DGE analysis in NUAK1-deficient cortical neurons.

### Nuclear speckle protein SON is a novel substrate of NUAK1

To identify how it can regulate neuronal RNA splicing, we performed an unbiased phospho-proteomic experiment to identify direct substrates of NUAK1, using an ATP analog-sensitive (AS) kinase assay (*42–44*) coupled with mass spectrometry (**Fig. 2A**). For this, methionine 133 in the catalytic site of NUAK1 was mutated into a glycine (hereafter M133G), to allow for the accommodation of a bulky ATP (6-benzyl-ATPγS). Transfer of the thiol-modified gamma phosphate enables immunolabelling of proteins directly phosphorylated by NUAK1. We validated that the AS-mutated, but not the wild-type NUAK1 could use the bulky ATP (**Suppl. Fig. 2A**) and validated the method by observing thio-phosphorylation of MYPT1, a well-characterized substrate of NUAK1 (*45*) (**Suppl. Fig. 2B**). Differential mass spectrometric analysis of SILAC labelled immune-precipitates from U2OS cells overexpressing either NUAK1^M133G^ or wild-type NUAK1 incubated with 6-benzyl ATPgS identified 3 high confidence (FDR<1%) putative substrates of NUAK1 (**Fig. 2B**): PPP1R12A (also known as MYPT1), isoleucyl-tRNA synthetase 1 (IARS), and the nuclear speckle protein SON.

**Figure 2.**
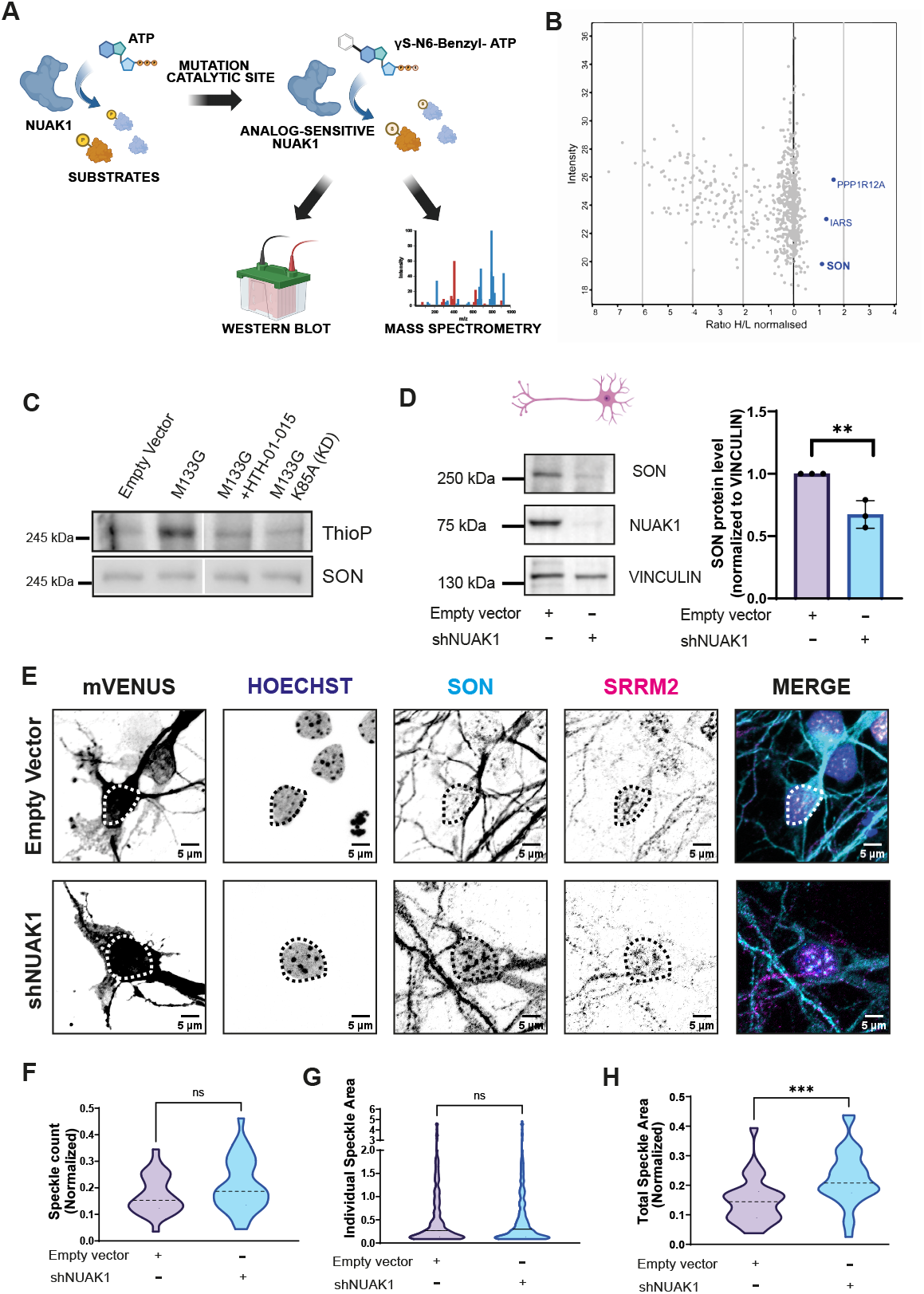
Nuclear speckle protein SON is a novel substrate of NUAK1. (A) Schematic representation of analog sensitive kinase assay workflow for proteomics or Western blot. (B) Plot showing proteins that were significantly phosphorylated by analog sensitive NUAK1 during mass spectrometry analysis. (C) Western blot representing the analog sensitive kinase assay with Flag-NUAK1^M133G^ detecting the phosphorylation of protein SON. (D) Western blot representing protein levels of SON in primary cortical neurons transduced with a control vector or shRNA against *Nuak1*, Paired t-test, *\*\* p < 0*.*01*. (E) Immuno-fluorescence assay showing nuclear speckles through staining of SON and SRRM2 in primary cortical neurons (5DIV). (F-H) Nuclear speckle parameters based on SRRM2 immuno-fluorescence in primary cortical neurons (5DIV): speckle count (F; normalized to nuclear area), individual speckle area (G), and total speckle area (H; normalized to nuclear area), Mann-Whitney test, *\*\*\* p < 0*.*001*.

To confirm that SON is a bona fide substrate of NUAK1, we performed *in vitro* kinase assays using the AS mutant protein (**Fig. 2C**). Here we observed a phosphorylation of SON that was lost by using the NUAK1-specific inhibitor HTH-01-015 (*46*) or by expressing a catalytically-inactive mutant protein (K85A). To confirm these results, we sought which functional domain of SON is phosphorylated by NUAK1. SON is a large protein, consisting of multiple functional domains: the N-terminus, followed by a repeat region, a putative DNA-binding domain, an arginine-serine rich (RS) domain and two RNA-binding domains (G-patch and double-strand RNA-binding motif) (**Suppl. Fig. 2C**) (*47*). Our mass spectrometry analysis indicated that NUAK1 is likely to phosphorylate residues in the C-terminal end of SON protein (data not shown), which contains the RS domain and two RNA-binding domains. We overexpressed HA-tagged human SON C-terminus (a.a. 1819-2426) in 293T cells and performed immuno-precipitation using an anti-HA antibody, followed by an *in vitro* kinase assay using AS-NUAK1-containing lysate (**Suppl. Fig. 2C-D**). We observed a clear band that is present at the height of the immune-precipitated C-terminus with the thioester-detecting antibody, while the band is absent in the empty vector condition. From this observation, we conclude that NUAK1 is able to phosphorylate the C-terminus of SON *in vitro*.

We next investigated the functional effect of NUAK1-mediated phosphorylation on SON. The inhibition of NUAK1 catalytic activity using HTH-01-015 (*46*) or shRNA-mediated knockdown of *Nuak1*, led to loss of SON protein levels, both in 293T cells and primary neurons, respectively (**Fig. 2D** and **Suppl. Fig. 2E**). Of note, quantitative PCR analysis demonstrated that *Son* mRNA levels were not altered in either condition, thus suggesting that NUAK1 regulates SON at the post-transcriptional level (**Suppl. Fig. 2F-G**).

Finally, we tested whether NUAK1 inhibition and the subsequent loss of SON would impact nuclear speckle structural features. Analysing the nuclear speckles through SRRM2 yielded a significant reduction in individual nuclear speckle area accompanied by a significant increase in nuclear speckle number after 72h of HTH-01-015 treatment in 293T cells (**Suppl. Fig. 2H-I-J**), without affecting the total coverage of the nuclear area by nuclear speckles (**Suppl. Fig. 2H-K**). Of note, NUAK1 inhibition led to an enlargement of the nuclei in treated cells (**Suppl. Fig. 2H-L**). We performed the same experiment in neurons, but since HTH-01-015 is lethal in primary cortical neurons, we used a shRNA-mediated knockdown of *Nuak1* (**Fig. 2E**). Similar to 293T cells, we observed an increased number of nuclear speckles, although the result was not statistically significant (**Fig. 2F**). Contrary to 293T cells, there was no change in individual speckle size (**Fig. 2G**), which translated into an increased total nuclear area occupied by nuclear speckles (**Fig. 2H**). Altogether, our data demonstrate that SON is a novel substrate of NUAK1. Downregulation of NUAK1 protein or reduction of its catalytic activity reduces SON protein levels, impacting nuclear speckle structural features both in dividing cells and post-mitotic neurons.

### NUAK1 and SON share a common splicing signature in neurons

To determine to what extent SON regulation contributes to the NUAK1-associated splicing signature in developing cortical neurons, we performed another RNA-sequencing experiment in DIV5 neurons in which *Son* was knocked down using lentiviral transduction. Analysis of the splicing dataset using rMATS revealed that knockdown of *Son* differentially affects 2845 splicing events (**Suppl. Fig. 3A**). The distribution of splicing events across the *Son* knockdown dataset was found to be similar to *Nuak1* knockdown: the majority of splicing events affected cassette exons (ES: 57.2%; MXE: 25.2%), while intron retention made up approximately 10% of the total splicing events (**Suppl. Fig. 3B)**. Like for *Nuak1, Son* knockdown equally biased the splicing towards more intron retention but, caused both exon skipping and inclusion at comparable levels (**Suppl. Fig. 3C-D**) and affected around 70% exons in the coding sequence of the protein (**Suppl. Fig. 3E**). *Son* knockdown impacted the splicing of genes primarily involved in fundamental neuronal functions, such as synaptic processes (regulation of synapse organization, vesicle-mediated transport of synaptic proteins), dendritic morphogenesis but also axon development (**Suppl. Fig. 3F**). This suggests that SON-mediated splicing could impact neuronal development and/or function. A differential gene expression analysis for *Son* knockdown yielded 1750 genes whose transcription was either upregulated (843 genes; 48.2%) or downregulated (907 genes; 51.8%) (**Suppl. Fig. 3G**). Like for NUAK1-deficiency, the genes that are present in the DGE analysis and the splicing analysis show minimal overlap (**Suppl. Fig. 3H**).

Next, we assessed the overlap between the *Nuak1* and *Son* knockdown splicing dataset. We found that 330 splicing events were shared between the two splicing datasets, i.e. the genetic features with the same chromosomal coordinates were affected with the same directionality (ΔPSI) (**Fig. 3A** and **Suppl. Fig. 4A**). Those splicing events retain a very similar repartition across splicing event type compared to the *Nuak1-* and *Son-* associated splicing datasets (**Fig. 3B**). In addition, we observed a positive correlation between the magnitude of the NUAK1-mediated splicing events and those of SON (**Fig. 3C**). The overlap in the splicing signature is in stark contrast with the lack of overlap between NUAK1 and SON datasets at the differential gene expression level (**Suppl. Fig. 4B**). The observation that the majority of common splicing events share the same directionality suggests that NUAK1 is a positive regulator of SON function. This is consistent with our observations about NUAK1 being important for the maintenance of SON protein levels.

**Figure 3.**
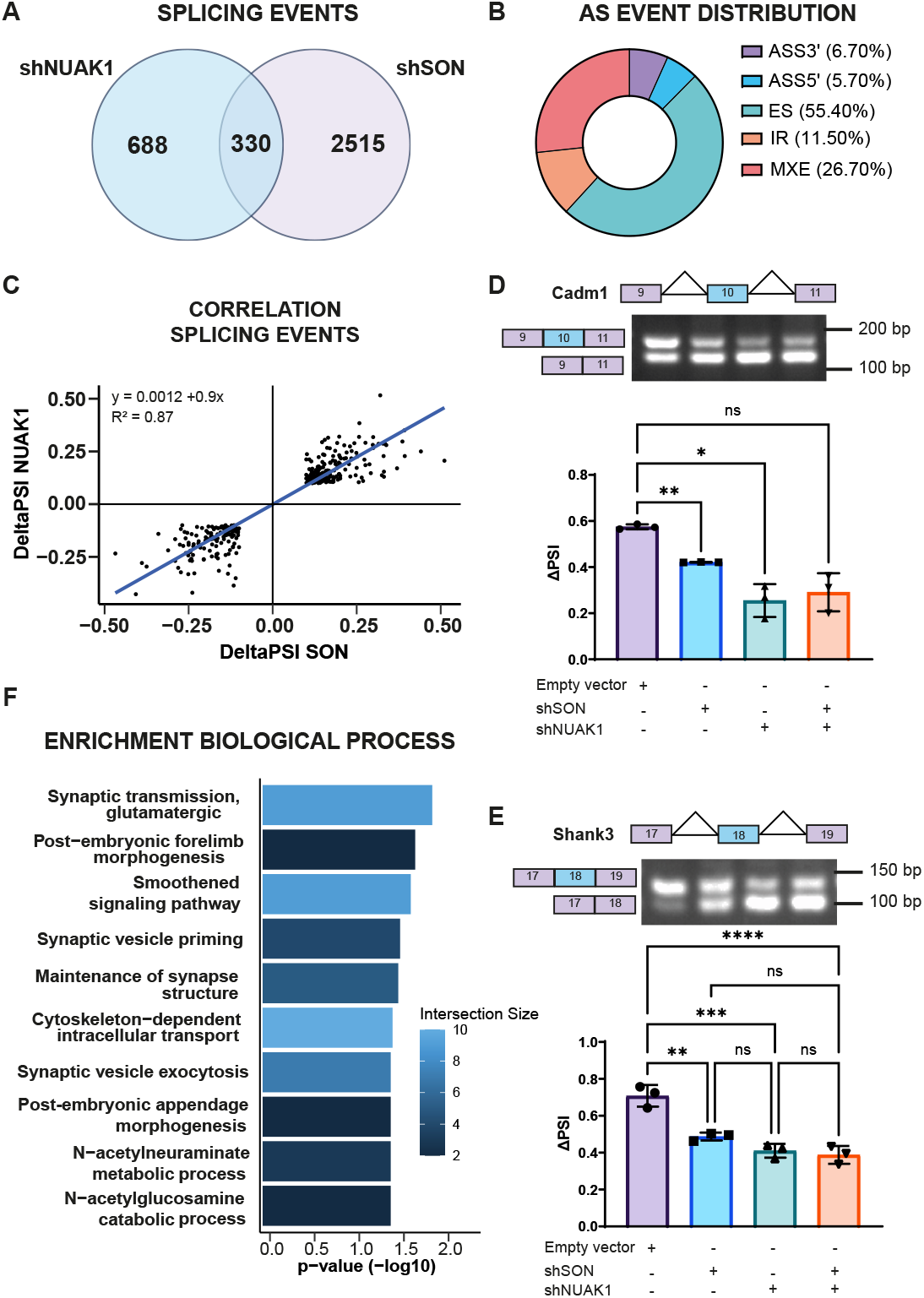
NUAK1 and SON share a common splicing signature in neurons. ((A) Overlap in genes affected by differential splicing between NUAK1- and SON-deficient cortical neurons (5DIV). (B) AS event distribution per category in the shared splicing signature between NUAK1- and SON-deficient neurons. (C) Scatter plot showing the ΔPSI of splicing events in NUAK1-deficient cortical neurons *versus* the ΔPSI of the same splicing event in SON-deficient cortical neurons. (D-E) PCR showing a splicing event affecting exon 10 in the *Cadm1* gene and exon 18 in the *Shank3* gene in cortical neurons (5DIV). Quantification of splice isoforms in the *Cadm1* and *Shank3* gene in various conditions of NUAK1- and SON-deficiency in cortical neurons (5DIV), RM one-way ANOVA and One-way ANOVA, respectively, *\* p < 0*.*05, ** p < 0*.*01, *** p < 0*.*001, **** p < 0*.*0001*. (F) gProfiler enrichment analysis for biological process on genes that were differentially spliced in both NUAK1- and SON-deficient primary neurons.

We validated some of the events from the shared splicing dataset using PCR to assess splicing differences in a semi-quantitative way (**Fig. 3D-E**). Our experiments show that knockdown of either *Nuak1* or *Son* significantly altered the inclusion of exon 10 in the *Cadm1* transcript in cortical neurons, as was observed from the RNASeq dataset. A second example is the inclusion of exon 18 in *Shank3* transcript, which is impaired upon knockdown of either *Nuak1* or *Son* (**Fig. 3E**). Simultaneous knockdown of *Nuak1* and *Son* did not have an additive effect on the splicing of *Cadm1* or *Shank3* transcripts, providing further evidence that NUAK1 and SON work in sequence in a common pathway (**Fig. 3D-E**). Conversely, we ruled out that the overlap between the NUAK1 and SON targets result from a general splicing blockage. Indeed, some splicing events were deregulated specifically upon knockdown of *Nuak1* but unchanged in the absence of *Son* (**Suppl. Fig. 4C-G**; *Ppiaf1* and *Stx3*). Conversely, we identified splice events altered by *Son* inhibition that were unchanged in the absence of *Nuak1* (**Suppl. Fig. 4C-G**; *Shank1* and *Prkcg*).

Lastly, we performed an enrichment analysis on the shared splicing dataset showing that the affected genes are mainly enriched in terms related to synaptic processes (e.g. maintenance of synapse structure, synaptic vesicle priming…) (**Fig. 3F**).

We conclude that NUAK1 and SON have an overlapping splicing signature that provides evidence for a functional interaction within a common regulatory pathway for splicing in cortical neurons.

### The shared splicing signature of NUAK1 and SON is enriched in highly conserved splicing events and genes associated with neurodevelopmental disorders

To test the relevance of the shared splicing signature between NUAK1 and SON, we tested for the enrichment in genes linked to neurodevelopmental diseases (NDDs) like epilepsy or autism spectrum disorder (ASD), or to neuroanatomical features such as head circumference (**Fig. 4A-B**). Yet, there seems to be a particular enrichment of the shared signature genes in the category of high-risk genes for NDDs and genes found in the SFARI database compared to both the *Son* and *Nuak1* splicing datasets (**Fig. 4A**). For example, we found a number of genes that are highly associated with ASD in the shared splicing signature between *Nuak1* and *Son*. These include *Shank3, Scn8a, Cadm1, Chd2, Auts2* and *Gria2*. This is underlined by an odds ratio analysis for the shared signature regarding several categories of NDDs (**Fig. 4B**).

**Figure 4.**
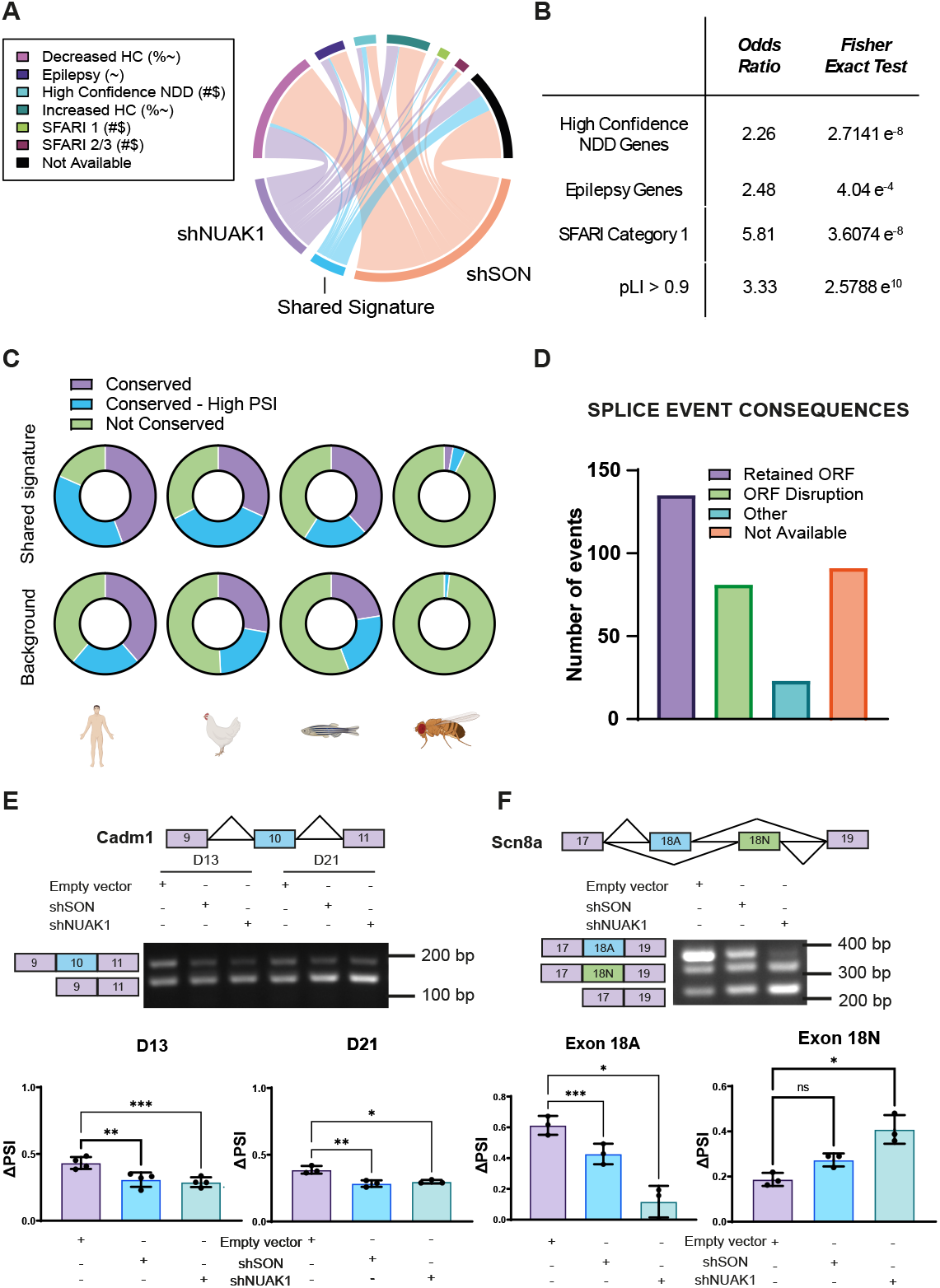
The shared splicing signature of NUAK1 and SON is enriched in highly conserved splicing events and genes associated with neurodevelopmental disorders. (A). Circlize plot showing the links between the genes affected by differential splicing in NUAK1- or SON-deficient cortical neurons, as well as the shared splicing signature and categories of neurodevelopmental disorders, as defined in the GeneTrek database. Hypergeometric test, # overrepresentation compared to NUAK1 dataset, $ overrepresentation compared to SON dataset, % underrepresentation compared to SON dataset,~ underrepresentation compared to NUAK1 dataset. The width of the links represents the relative proportion between the three datasets. (B) Odds ratio and its statistical significance for gene enrichment of the shared splicing signature in various classes of neurodevelopmental diseases, as classified by the GeneTrek database. (C) Conservation of splicing events (as defined in the VastDB database) in the shared splicing signature between NUAK1- and SON-deficient cortical neurons compared to the background dataset of expressed genes in cortical neurons. (D) Impact of the splicing events in the shared signature at the protein level as predicted by VastDB. (E) Developmental splicing pattern across 13DIV and 21 DIV of exon 10 in *Cadm1*, RM One-way ANOVA, *\* p < 0*.*05, ** p < 0*.*01, *** p < 0*.*001*. (F) Splicing pattern of exons 17-18(A/N)-19 of *Scn8a* in cortical neurons (5DIV) and its quantification, RM One-way ANOVA and Friedman test, respectively, *\* p < 0*.*05, *** p < 0*.*001*.

To gain a better insight in the impact of the NUAK1-SON pathway in splicing, we decided to take a look from an evolutionary angle. We cross-referenced our shared splicing events with VastDB, a database compiling information about splicing events across different tissues and species (*48*). 73% (239 out of 330) of the splicing events in the shared signature were present in the VastDB database (**Fig. 4C**). Generally, the conservation levels for splicing events in vertebrates were higher compared to those found in splicing events for the genes outside the shared signature in our RNASeq dataset. This was especially true for splicing events that showed high levels of inclusion. However, we found an important enrichment (approximately 5-fold) in conserved events in Drosophila, compared to the background dataset, suggesting that the NUAK1-SON shared splicing signature affects splicing events that are very highly conserved (**Fig. 4C**). At the protein level, VastDB predicts that an important number of splicing events shared between NUAK1 and SON (59.1%, 135 out of 239) would retain an open reading frame (ORF) and thus the splicing event would switch between protein isoforms (**Fig. 4D**). However, 34% (82 out of 239) of splicing events would cause a disruption of the ORF with unknown consequences. Next, we wanted to understand whether NUAK1 and SON are affecting very early neurodevelopmental splicing or whether later developmental stages would be impacted as well. It is well established that splicing patterns change during neurodevelopment and that this is strictly regulated in time and space (*49*). For example, an exon inclusion event in *Cadm1* splicing, was diminished in favour of the exon exclusion form during neuronal maturation (**Fig. 4E** and **Fig. 3D**). However, not only during early time points (DIV5), but also during dendritic development and synaptic maturation, deficiency of NUAK1 or SON caused skipping of the exon (**Fig. 4E**). We observed similar results for an exon inclusion event concerning exon 18 in *Shank3* mRNA (**Suppl. Fig. 4H**).

Lastly, we took a closer look at the 17 splicing events that were conserved in Drosophila. Among those events, we found an exon skipping event at exon 18 in the *Scn8a* transcript, encoding the sodium channel Nav1.6. This sodium channel is mainly expressed at the axon initial segment and nodes of Ranvier, and is crucial for the electrophysiological properties of excitatory neurons (*50*). The splicing event from *Scn8a* we found in our dataset consists of two alternative exons 18: exon 18A and exon 18N (*51, 52*).

Whereas exon 18A encodes the fully functional ion channel, exon 18N causes non-sense-mediate decay. Moreover, the inclusion of exon 18A and 18N is developmentally regulated in the brain (*51, 52*). Upon neuronal maturation, the exon 18A becomes the predominant form, over 18N (*51, 52*). According to our RNASeq dataset, SON and NUAK1 deficiency would induce skipping of exon 18A, which we confirmed using PCR (**Fig. 4F**). Inclusion of exon 18N, on the other hand, was increased. This shows that splicing changes induced by *Nuak1* or *Son* deficiency could have an important functional effect on neurons.

In all, these data provide evidence that NUAK1 and SON regulate a subset of highly conserved genes, that are crucial for neurodevelopment. Their function is not limited to one stage of development, but rather maintain neurodevelopmental splicing patterns throughout neuronal maturation.

### SON deficiency phenocopies NUAK1 deficiency with respect to neuronal morphology and maturation

Finally, we wondered to what extent SON deficiency would mimic a lack of NUAK1 in developing cortical neurons. We first characterized the impact of *Son* knockdown on axon development using murine primary cortical cultures and the *ex utero* electroporation technique (**Fig. 5A**). We electroporated neural progenitor cells with a plasmid encoding mVenus and either an empty vector (control) or a plasmid expressing an shRNA against SON (shSON). After 5 days of culture, we observed that cortical neurons lacking SON, had shorter axons and fewer branches (normalized for axon length) compared to control neurons (**Fig. 5B-C**). This phenotype closely resembles what was observed upon NUAK1 deficiency (*25–27*).

**Figure 5.**
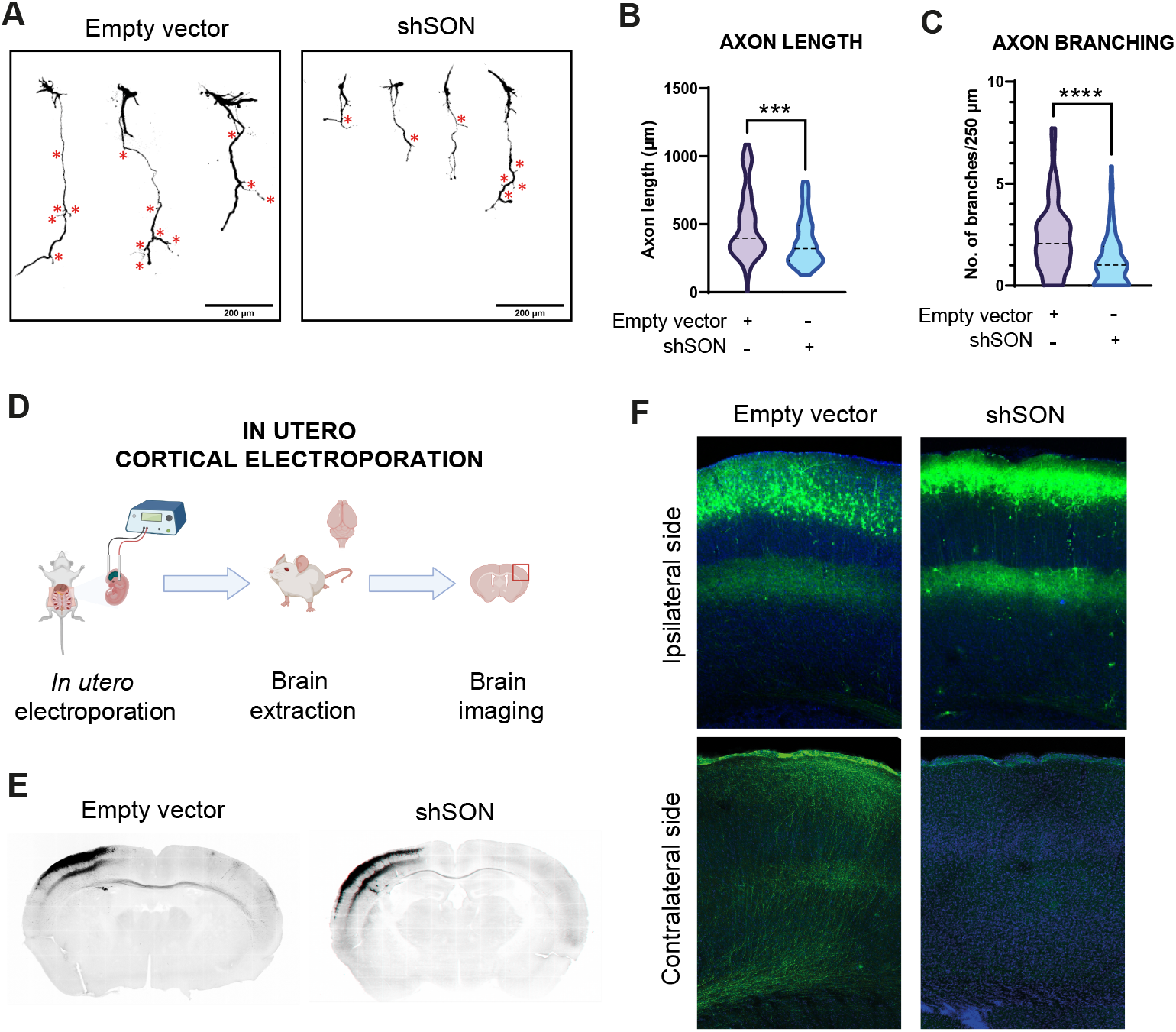
SON deficiency partially phenocopies NUAK1 deficiency in terms of neuronal morphology and maturation. (A) Representative images of primary cortical neurons at 5DIV. (B-C) Axon length and branching quantification in early developing cortical neurons (5DIV) in the presence or absence of SON, Mann-Whitney test, *\*\*\*\* p < 0*.*0001*. (D) Schematic representation of *in utero* electroporation in primary cortical neurons. (E) Typical whole brain slice after *in utero* electroporation with pSCV2 and control plasmid or a plasmid encoding shSON. GFP signal represented in black. (F) Detail on the electroported (ipsilateral; P21) side showing neuronal cell bodies and short-range axonal connectivity, and long-range projections (contralateral projections; p21);

Secondly, we decided to look at dendritic complexity *in vitro* at 13DIV. SON-deficient neurons showed a marked reduction in dendritic complexity, as analysed by using the Sholl method (**Suppl. Fig. 5A-B**). This reduction in dendrites is mostly situated close to the cell body.

Lastly, we looked at the effect of *Son* knockdown in an *in vivo* setting by performing *in utero* electroporation on pregnant dams at E15.5 (*33*) (**Fig. 5D**). The embryonic somatosensory cortex was electroporated with a similar mix of plasmids as for the *ex utero* electroporation. At P21, electroporated animals were sacrificed and brains were processed to visualize neuronal development and cortical circuits formation (**Fig. 5E**). On the ipsilateral side, we observed a small but reproducible effect of SON inhibition on neuronal migration, similar to previous observations (*53*) (**Fig. 5F**). Furthermore, dendritic development was impaired, with an apparent reduction of the apical dendrite. Despite the presence of axons in reaching the white matter and the corpus callosum, we saw a strong reduction of long-range projections to the contralateral somatosensory cortex (**Fig. 5F**). From this set of experiments, we conclude that SON deficiency critically impacts neuronal morphological development both *in vitro* and *in vivo*. Thereby, lack of SON partially phenocopies a lack of NUAK1 in cortical neurons, but simultaneously, SON knockdown displays a distinct phenotype for dendritic complexity.

## DISCUSSION

In this study, we present a novel mechanism through which alternative splicing is regulated in developing cortical neurons, with NUAK1 and SON as key players. We propose that SON is a novel, direct effector of the serine/threonine kinase NUAK1 in cortical development. This is supported by evidence that 1) NUAK1 and SON co-localize to nuclear speckles, 2) NUAK1 phosphorylation is necessary for the maintenance of SON at the protein level, ultimately impacting nuclear speckles, 3) together, NUAK1 and SON regulate a subset of strongly conserved splicing events, impacting neuronal development, 4) NUAK1 and SON demonstrate epistasis on the regulation of mRNA splicing, and 5) *Son* knockdown phenocopies the axonal phenotypes previously associated with NUAK1. More broadly, we also provide compelling evidence for a role of SON as a regulator of neurodevelopment, which sheds light to potential neuronal alterations in ZTTK syndrome. Altogether, *Son* deficiency impacts a number of critical processes during neurodevelopment, notably axon growth and branching, and dendritic development both *in vitro* and *in vivo*.

### NUAK1 is a critical regulator of RNA splicing in developing cortical neurons

NUAK1, a kinase whose mutations are associated with various NDDs (*54–57*), as well as cerebral malformations (*58*), was previously characterized as a regulator of axon outgrowth and branching in the developing mouse cortex (*25, 26*). The initial evidence pointed to altered mitochondrial distribution in the axon and impaired metabolism (*27*). Here, we demonstrate an alternative mechanism by which NUAK1 impacts neuronal development, through the regulation of neuronal splicing. Analyses of the splicing signature associated to *Nuak1* knockdown failed to uncover metabolic pathways, suggesting that splicing and metabolic regulation are two distinct functions of NUAK1 in neurons. How these two seemingly distinct functions of NUAK1 are cross-regulated remains unknown. As a matter of fact, evidence points to a crosstalk between metabolic regulation and alternative splicing. Interestingly, the Arginyl tRNA synthetase RARS can act as a sensor of cellular arginine level to regulate the nuclear localization of speckle protein SRRM2 (*59*). Given that our proteomic analysis suggests that NUAK1 can phosphorylate IARS, one can speculate that NUAK1 may act as a global coordinator of metabolic level, protein synthesis, and alternative splicing in developing neurons.

Our transcriptomic dataset showed 823 genes in which splicing was significantly affected by knocking down *Nuak1*. This is almost sixfold the number of genes that were impacted at the gene expression level. Moreover, the genes that are differentially expressed in the *Nuak1* knockdown condition are not the same genes that are affected by the differential splicing. One possible explanation for this discrepancy might be that NUAK1 would act at several levels on the gene expression program, regulating independently transcription itself and co- or post-transcriptional processes. However, it is equally likely that network effects and trans-regulation of RNA-binding proteins could indirectly lead to changes in transcription or RNA stability. Although the same reasoning could be applied to alternative splicing, our data rule out a general deregulation of the splicing machinery, as evidenced by the fact that some exons were specific from NUAK1 and not affected by the downregulation of *Son*, and conversely *Nuak1* knockdown didn’t affect some exons regulated by SON.

### Nuclear speckle protein SON is a novel effector of NUAK1

In accordance with a role for NUAK1 in the regulation of neuronal splicing, we show that NUAK1 localizes to the nucleus and associates with nuclear speckles, matching previous observations that NUAK1 colocalizes with nuclear speckle proteins in cancer cells (*28*). Recent evidence has emerged for a proline-rich sequence that targets proteins to the nuclear speckles and that NUAK1 harbours such a sequence (*60*). Interestingly, such sequence is also found in the close paralog NUAK2, but not shared with any other constituent of the AMPK-related family of kinases, consolidating the notion that spliceosome regulation is a core molecular function of the NUAK proteins. Nonetheless, it may be interesting to investigate how the various AMPK-related kinases works together and whether their respective signalling pathways intersect to influence splicing. One could for example imagine an interplay with AMPK, which has recently been shown to phosphorylate SRSF1 to impact splicing (*61*).

Although the precise mapping of the NUAK1 target site was impossible owing to the arginine and serine rich nature of SON, we could narrow down that the C-terminus of SON is being phosphorylated by NUAK1, without ruling out that other parts of the protein might be phosphorylated by NUAK1 as well. The C-terminus end of SON contains a RS domain that is present in many splicing factors and which is believed to regulate their interactions (*62, 63*), but could also promote nuclear localization of splicing factors through the SR-protein nuclear import receptor, transportin-SR (*64, 65*). One hypothesis could therefore be that phosphorylation by NUAK1 may change the interactome of SON, thereby impacting splicing. Furthermore, the phosphorylation by NUAK1 seemingly impacts the stability of SON, given that long-term knockdown or inhibition of NUAK1’s kinase function leads to a loss of SON at the protein level. SON is a ubiquitously expressed protein, present in the majority of tissues (*66*). Despite this common expression pattern, it seems that SON is able to target distinctly specific neuronal genes and splicing events in developing neurons. A possible explanation could be that SON by itself does not harbour a specific preference for certain splice sites, but that it gets its specificity from interacting with other splicing factors that are tissue specific or cell type-specific. For example, recent work has shown that SON and RBFOX2 compete with each other for binding of RNA in glioblastoma cells (*67*). Given the conservation of the RNA recognition motif between RBFOX1, −2 and −3 (*68–70*), it is likely that all of them would functionally interact with SON. However, RBFOX1, −2, and −3 have distinct tissue expression patterns (*68–70*), which could render the functional interaction with SON and thus the resultant splicing, tissue-specific. This specificity may be additionally finetuned by interaction with signalling pathways and NUAK1, could be one such regulator of the SON interactome to regulate SON’s splicing properties. Interestingly, NUAK1 is strongly expressed in the developing telencephalon, with a peak expression during the earliest stages of cortical development. Thus, the phosphorylation of SON by NUAK1 might be a way to spatially or temporally regulate a SON-dependent alternative splicing program in the developing nervous system. In general, it remains to be resolved how SON is embedded in the splicing regulatory network, not only in neurons but also in other tissues.

### The NUAK1-SON pathway regulates a highly conserved subset of splicing events in neurodevelopment

From the overlap between the NUAK1 and SON RNASeq dataset, we conclude that NUAK1 is a positive regulator of SON and that NUAK1’s effect on splicing is for an important part mediated by SON. Interestingly, there is an enrichment of genes connected to NDDs in the shared subset and it contains a number of splicing events that are highly conserved. This suggests that both NUAK1 and SON would regulate neurodevelopment up and down the evolutionary tree. Therefore, it may be interesting to study their interaction in less complex models like *Drosophila*. Both NUAK1 and SON orthologs are present in *Drosophila*. For SON it is the C-terminus with the RS- and RNA-binding domains that is conserved, but more fascinatingly, *Drosophila* SON is only 843aa long, compared to its 2426aa long human counterpart. This opens up the question what constitutes the function of the unique N-terminal repeat-domains that are present in mammals and what would be their impact on splicing and nuclear speckle biology in general.

This impact on splicing during neurodevelopment and specifically the impact on highly conserved splicing events, may explain the effects that we see on neuronal morphology *in vitro* and long-range projections *in vivo*. These findings are consistent with the neurodevelopmental aspect of patients with the Zhu-Tokita-Takenouchi-Kim (ZTTK) syndrome. This is a broad-spectrum developmental syndrome which is caused by heterozygous SON loss-of-function mutations and affects multiple organs, but notably the central nervous system (*17, 30-32, 71*). Patients display brain malformations, epilepsy, ASD-like behaviour and intellectual disability, but also facial dysmorphia and general affectation of the musculoskeletal system. The particular severity of SON deficiency in these organ systems with an observable impact early on in infancy and childhood may be related to their high reliance on spatiotemporal control of alternative splicing (*49*). Understanding the exact role of SON and its interaction partners in not only splicing itself but its developmental regulation may provide a stepping stone towards understanding the pathology of SON deficiency.

### Conclusions

We provide evidence for a novel pathway to control splicing in cortical neurons involving the kinase NUAK1 and the splicing co-factor SON. They regulate a highly conserved subset of splicing events crucial for neurodevelopment. In addition, SON itself seems to be indispensable for proper development and maturation of cortical neurons. This role provokes its own questions and merits a line of research by itself.

## MATERIALS AND METHODS

### Animals

Experiments were carried out with respect to the French legislation regarding animal experimentation. Experimental protocols were approved by the ACCeS Ethics committee and was authorized by the French ministry of higher education and research. Animals were allowed *ad libitum* access to food, water and were maintained on a 12-hr light-dark cycle. For cell culture, pregnant Swiss females were purchased from Janvier Labs. They were transported to the animal facility at E13.5 (embryonic day). For the IUE, timed-pregnant females hybrid F1 (C57Bl/6Jrj x 129Sv) were obtained by overnight breeding with male C57Bl/6Jrj. Noon following breeding was considered as E0.5. The experiments were performed at E15.5.

### Cell culture and transfection of cell lines

293T cells were cultured at 37°C, 5% CO_2_ in Dulbecco’s Modified Eagle’s Medium (DMEM), supplemented with 10% foetal bovine serum (FBS). For transfection purposes, cells were seeded as such to reach 50 % confluence on the day of transfection. Cells were transfected with jetPrime (#101000015, Polyplus) according to the manufacturer’s instructions. After 4h, the medium was exchanged for fresh medium.

### Neuronal cell culture and electroporation

The electroporation of dorsal telencephalic progenitors was performed by injecting plasmid DNA (1-2 μg/μL of endotoxin-free plasmid DNA) plus 0.5% Fast Green (Sigma; 1:20 ratio) using a Picospritzer III microinjector (Harvard Apparatus) into the lateral ventricles of isolated E15.5 embryonic mouse heads. Electroporations were performed with gold-coated electrodes (GenePads 5 mm, BTX) using an ECM 830 electroporator (BTX) and the following parameters: five pulses of 100 milliseconds (msec), 150 msec interval, at 20 V. Immediately after electroporation, cortices were dissected in Hank’s Buffered Salt Solution (HBSS) supplemented with HEPES (pH 7.4; 2.5 mM), CaCl_2_ (1 mM, Sigma), MgSO_4_ (1 mM, Sigma), NaHCO_3_ (4mM, Sigma) and D-glucose (30 mM, Sigma), hereafter referred to as cHBSS. Cortices were dissociated in cHBSS containing papain (10 U/mL, Worthington) and DNAse I (100 μg/ml, Sigma) for 15 min at 37 °C, and washed three times. Cells were then counted and seeded at an appropriate density in plates coated with poly-D-lysine (1 mg/ml, Sigma) and laminine (0.01mg/mL). Cells were cultured for 5-21 days in Neurobasal medium supplemented with B27 (1X), N2 (1x), L-glutamine (2 mM) and penicillin (5 U/mL)-streptomycin (50 mg/mL). For neuronal cultures lasting longer than 5 days, 5% FBS was added and starting from DIV5, half of the medium was changed every other day.

### In Utero Cortical Electroporation and brain sectioning

In Utero Cortical Electroporation were performed at E15.5 as described in (*33*). A mix containing 1 μg/μl endotoxin-free plasmid DNA plus 0.5% Fast Green (Sigma; 1:20 ratio) was injected into one lateral hemisphere. Electroporation was performed with platinum electrodes (Tweezertrodes 3 mm, BTX Harvard Apparatus, ref:45-0487) using an ECM 830 electroporator (BTX) and the following parameters: four pulses of 100 msec, 500 msec interval, at 45V. Animals were sacrificed at Postnatal day 21 by Intracardiac perfusion of 1x PBS (total volume 5mL) and then terminal perfusion of 4% paraformaldehyde/1X PBS (PFA, Electron Microscopy Sciences) followed by 24h post-fixation in 4% PFA. After replacing the PFA with PBS, brains were embedded in 3% low-melting point agarose (MP Biomedicals; #AGAL0050) and sectioned (thickness: 80 µm) with a vibratome (Leica VT1000S). Afterwards, sections were immuno-stained as described below.

### Immuno-precipitation and kinase assay

293T cells were transfected with SON C-terminus, pCIG2-Flag-NUAK1^M133G^ or pCIG2-Flag-NUAK1^M133G/K85M^. SON-overexpressing cells were washed with PBS 1x and lysed after 48h using NP40 lysis buffer (*cfr. supra*). Cell lysates were collected and 200 µL of lysate was incubated overnight (approximately 16h) with anti-HA primary antibody at 4°C on a rotor. The next day, Flag-NUAK1^M133G^-overexpressing cells were washed with PBS 1x and lysed in NP40 buffer and 200 µL of lysate was incubated with anti-Flag primary antibody for 4h at 4°C on a rotor. Afterwards, lysates were incubated with 50 µL of ProteinA-Sepharose™ 4 beads (P9424, Sigma) for 3h on a rotor. Before use, beads were washed two times with NP40 buffer. Subsequently, beads were collected and washed two times with kinase buffer (20 mM Tris-HCl pH 7.5, 10 mM MgCl_2_, protease inhibitor cocktail without EDTA (Roche, 11836170001), 1 mM DTT and 100 µM ATP). After washing beads from both lysates were added together in the same tube and incubated with 150 µL of kinase buffer, including 6-benzyl-ATPγS (500 µM, BioLog, B-072-05). This reaction mix was incubated for 1h at 30 °C under light agitation. The reaction was ended by adding EDTA (Promega, V4233) at a final concentration of 20 mM and incubated at room temperature during 10 min. Lastly, an alkylation reaction was initiated by adding p-nitrobenzylmesylate (PNBM, 5 mM final concentration) for 1h at room temperature. Laemmli buffer (4x, 200 mM Tris-HCl pH 6.8, 400 mM DTT, 8% sodium dodecyl sulphate (SDS), 6 mM bromophenol, 4.3M glycerol) was added to the reaction mixtures and they were heated for 2 min at 95°C for complete denaturation. Samples were run on a Western blot and detection of the thiophosphate mark was done using thiophosphate ester recombinant rabbit monoclonal antibody (SD2020, ThermoFisher).

### RNA isolation, PCR and qPCR

Cells were harvested in 500 µL TRIzol™ (ThermoFisher, 15596026) according to the manufacturer’s instructions. Subsequently, the RNA was extracted using 250 µL of chloroform and phase separation by centrifugation for 15 min at 12,000 *g*, 4°C. The aqueous phase was transferred to a new tube and isopropanol was added at a ratio 1:1. After 10 min incubation at 4 °C, the samples were centrifuged for 10 min at 12,000 *g*, 4°C. The supernatant was removed and RNA pellets were washed with 500 µL of 70% ethanol. After removal of the ethanol, RNA pellets were left to dry to the air for 10 min and subsequently an appropriate amount of nuclease-free H_2_O was added. RNA was incubated for 15 min at 65°C in a heat block, before being cooled down on ice. RNA concentration and purity were determined using the NanoDrop 2000 Spectrophotometer (ThermoScientific).

Reverse transcription was performed mixing 500 ng of RNA with 10 mM dNTPs (EuroMedex, EUA01601, final concentration 500 µM), OligodT 12-18 primer (Invitrogen, 18418012, final mass 25 µg), 10 µM DTT and 5x First Strand Buffer (SuperScript II Reverse transcriptase 200 U/µL, Invitrogen, 18064-014, final concentration 1x), reaching a final volume of 19 µL. SuperScript™ II reverse transcriptase (SuperScript II Reverse transcriptase 200 U/µL, Invitrogen, 18064-014) was added at 1 µL per reaction. The reaction mix was incubated for 50 min at 42°C and the enzymatic reaction was inactivated by placing the mix at 70°C for 15 min.

To perform semi-quantitative splicing assays, we used PCR. Following mix was made: 10x DreamTaq Green buffer (ThermoFisher, B71, final concentration 1x), 10 mM dNTPs (EuroMedex, EUA01601, final concentration 200 µM), forward primer (final concentration 1 µM), reverse primer (final concentration 1 µM), DreamTaq Polymerase 5U/µL (ThermoFisher, EP0701, final concentration 1.25U). To this reaction mix 12.5 ng of cDNA was added per reaction. The PCR reaction was performed using a C100 Touch™ Thermal Cycler (Biorad) as follows: 1. Initial denaturation at 98°C for 30”, with a subsequent denaturation step for 10”; 2. Annealing of the primers for 30” at 55°C and an elongation step at 72°C for another 30”. This cycle was repeated 43 times restarting at the second denaturation step. A final elongation step was done for 10’ at 72°C.

To determine SON expression levels, we used RT-qPCR. For this, a mastermix of FastStart Universal SYBRGreen 2x (Merck, 04913850001, final concentration 1x) and forward and reverse primer was made (1 µM final concentration for each). To every reaction, 30 ng of cDNA was added. RT-qPCR was performed using the CFX Connect™ Real-Time System (Biorad) as follows: 1. Initial denaturation at 95°C for 3’, with a subsequent denaturation step for 10”; 2. Annealing of the primers for 10” at 59°C and an elongation step at 72°C for 30”. This cycle was repeated 40 times restarting at the second denaturation step. A final elongation step was done for 10’ at 72°C.

### Immuno-fluorescence assay, image acquisition and image analysis

Cells were fixed for 30 min with 4 % PFA and washed three times with PBS 1x. Afterwards, cells were permeabilized using a permeabilization buffer (PB: PBS 1x, 0.3% Triton-X, 0.3% BSA) for 30 min. Cells were incubated afterwards with primary antibody diluted in PB for 1h at room temperature (anti-GFP) or overnight at 4 °C. After washing three times with fresh PB, Hoechst solution (1:10 000; Sigma; 94403) and secondary antibody diluted in PB (1:2000) were added and incubated with the cells for 1h at room temperature. Before mounting coverslips on glass slides with Fluoromount-G™ mounting medium (Invitrogen; 00-4958-02) for imaging, cells were washed three times with PBS 1x. Coverslips were imaged using a Nikon Eclipse Ti inverted microscope, equipped with an A1R detection system. Images were analyzed using Fiji software.

### Statistics

For every dataset, normality was determined and subsequently, the appropriate statistical testing was performed using Graphpad Prism 10 software. Statistical significance is indicated in all figures by the following annotations: ns, not significant; *, p<0.05; **, p<0.01; ***, p<0.001.

### RNASeq and downstream analysis

Primary cortical neurons were harvested at DIV5, 4 days after transducing the cells with control virus, a virus expressing an shRNA against SON or a virus expressing an shRNA against NUAK1. RNA was isolated using TRIzol™ in combination with RNA-binding columns. Library preparation consisted of removal of ribosomal RNA and ethanol precipitation. After fragmentation, first strand cDNA was synthesized using random hexamer primers. During second strand cDNA synthesis, dUTPs were replaced with dTTPs. The directional library was ready after end-repair, A-tailing, adapter ligation, size selection, USER enzyme digestion, amplification and purification. Samples were sequenced using Illumina paired end sequencing on an Illumina NovaSeq 6000, resulting in an average of 88301513 pairs of reads per sample. The quality control of the sequencing data was evaluated using FastQC (v0.11.9). The reads were trimmed using Prinseq-lite (v0.20.4) (*34*) (--trim-right 20) and filtered by average quality score (--trim-qual 20) and cutadapt (v4.1) (*35*). Reads were mapped EnsEMBL GRCm38.99 mouse reference using rna-STAR (v2.7.10a) (*36*). Reads below a mapping score of 10 or multimapped were filtered using samtools (v0.1.13) (*37*). The gene expression level in each sample was quantified with HTSeq-count (v0.13.5) (*38*). The differential gene expression (DEG) between conditions was calculated with DESeq2 (v1.38.0 using R v4.2.2) (*39*). We considered that genes were differentially expressed when their adjusted *p*-value was < 0.05, a baseMean ≥ 20 and |log2FoldChange| ≥ 0.4 (options: lfcThreshold = 0.4, altHypothesis = ‘greaterAbs’). Each condition was analysed using the Multivariate Analysis of Transcript Splicing (rMATS) program (*40*) to identify alternatively skipped exons, alternative 3′ splice sites, alternative 5′ splice sites, mutually exclusive exons, and retained introns. A filter was then applied on exon skipping events detected to select significant variants with an adjusted *p-*value ≤ and a ΔPSI ≥ 10%. Data are available on the GEO database (GSE294659).

Enrichment analysis was performed using gProfiler (update 2024-01-25) with Benjamini-Hochberg FDR (threshold = 0.05) against a background of genes attaining at least 10 reads in all samples of the control, shSON and shNUAK1 RNASeq dataset. To test the involvement of the differentially spliced genes in neurodevelopmental diseases, we cross-referenced those genes with the GeneTrek database. Odds ratios and significance were calculated using a tool on the GeneTrek database website (https://genetrek.pasteur.fr/). Evolutionary conservation and splicing event consequences were estimated by cross-referencing our dataset with VastDB (https://vastdb.crg.eu), an online database for alternative splicing events. The background dataset was the same dataset used for the gProfiler analysis. Lastly, for analysis of the position of splicing events (5’UTR, CDS or 3’UTR), data were converted from GRCm38 to GRCm39, using the Ensembl Assembly Converter. Subsequently, the relevant genes and chromosomal coordinates (GRCm39) were extracted from Ensembl using BioMart and our datasets were cross-referenced.

## Supporting information

Supplementary table 1: background genes

Supplementary table 2: shared signature

Supplementary table 3: NUAK1 signature

Supplementary table 4: SON signature

## ACKNOWLEDGEMENTS

The authors are grateful to members of the Courchet team for scientific discussions and suggestions. We thank Dr Franck Polleux (Columbia University, New York) and Dietmar Schmucker (LIMES institute, Bonn) for critical insight and feedback on the manuscript. We acknowledge support from the PSMN (Pôle Scientifique de Modélisation Numérique) of the ENS of Lyon for computing resources. We thank the personnel from the SCAR and ALECS-SPF mouse facility for animal care. This work was supported by the AFM-telethon through the strategic MyoNeurALP alliance and an ERC Starting Grant (678302-NEUROMET, J.C.). This work was performed within the framework of the LABEX CORTEX (ANR-11-LABX-0042 / ANR-11-IDEX-0007, J.C.). M.K. was the recipient of a postdoctoral fellowship from the Fondation pour la Recherche Medicale (SPF202110014126) and a Marie Sklodowska Curie Action fellowship (101110819).

## AUTHORS CONTRIBUTION

Authors contribution is listed below according to the CRediT (Contributor Roles Taxonomy) guidelines: **MK**: conceptualization, validation, formal analysis, investigation, data curation, writing original draft, visualization. **GMD**: Investigation, visualization, data curation. **HP**: Software, formal analysis, data curation. **SY**: Investigation. **JTM**: Methodology, investigation. **SL**: Formal analysis, data curation, visualization. **DS**: Formal analysis, data curation, visualization. **DJM**: Resources, visualization. **EYEA**: Resources, CFB: Resources, conceptualization. conceptualization. **CFB**: **EG**: Resources, Conceptualization, investigation, validation, formal analysis, data curation. **JC**: Conceptualization, writing original draft, funding acquisition, supervision, project administration. All authors have contributed to editing and reviewing the manuscript.

## REFERENCES

1. Z. Molnár., New insights into the development of the human cerebral cortex. journal of Anatomy 235, (2019/09/01).

2. P. Abe., Molecular programs guiding arealization of descending cortical pathways. Nature 2024 634:8034 634, (2024-09-11).

3. E. Klingler., Temporal controls over inter-areal cortical projection neuron fate diversity. Nature 2021 599:7885 599, (2021-11-09).

4. R. L. Walkerl., Genetic Control of Expression and Splicing in Developing Human Brain Informs Disease Mechanisms. Cell 179, (2019/10/17).

5. X. Qianl., Spatial transcriptomics reveals human cortical layer and area specification. Nature 2025 644:8075 644, (2025-05-14).

6. J. A. Calarco., Regulation of Vertebrate Nervous System Alternative Splicing and Development by an SR-Related Protein. Cell 138, (2009/09/04).

7. P. L. Boutzl., A post-transcriptional regulatory switch in polypyrimidine tract-binding proteins reprograms alternative splicing in developing neurons. Genes & Development 21, (2007-07-01).

8. B. Raj, Benjamin J. Blencowe, Alternative Splicing in the Mammalian Nervous System: Recent Insights into Mechanisms and Functional Roles. Neuron 87, (2015/07/01).

9. P. Mazinl., Widespread splicing changes in human brain development and aging. Molecular Systems Biology 9, (2013-01-22).

10. C. K. Vuong., The neurogenetics of alternative splicing. Nature Reviews Neuroscience 2016 17:5 17, (2016-04-20).

11. E. Furlanis, P. Scheiffele, E. Furlanis, P. Scheiffele, Regulation of Neuronal Differentiation, Function, and Plasticity by Alternative Splicing. Annual Review of Cell and Developmental Biology 34, (2018/10/06).

12. A. Joglekarl., Single-cell long-read sequencing-based mapping reveals specialized splicing patterns in developing and adult mouse and human brain. Nature Neuroscience 2024 27:6 27, (2024-04-09).

13. E. Furlanis., Landscape of ribosome-engaged transcript isoforms reveals extensive neuronal-cell-class-specific alternative splicing programs. Nature Neuroscience 2019 22:10 22, (2019-08-26).

14. Y. Zhang et., An RNA-Sequencing Transcriptome and Splicing Database of Glia, Neurons, and Vascular Cells of the Cerebral Cortex. Journal of Neuroscience 34, (2014-09-03).

15. S. M. Weyn-Vanhentenryck., HITS-CLIP and integrative modeling define the Rbfox splicing-regulatory network linked to brain development and autism. Cell reports 6, (2014 Mar 6).

16. S. Cuinat., Loss-of-function variants in SRRM2 cause a neurodevelopmental disorder. Genetics in Medicine 24, (2022/08/01).

17. J.-H. Kim, De Novo Mutations in SON Disrupt RNA Splicing of Genes Essential for Brain Development and Metabolism, Causing an Intellectual-Disability Syndrome. The American Journal of Human Genetics 99, (2016/09/01).

18. D. Li., Spliceosome malfunction causes neurodevelopmental disorders with overlapping features. The Journal of Clinical Investigation 134, (2024/01/02).

19. D. Greene., Mutations in the U4 snRNA gene RNU4-2 cause one of the most prevalent monogenic neurodevelopmental disorders. Nature Medicine 30, (2024 May 31).

20. Y. Chen., De novo variants in the RNU4-2 snRNA cause a frequent neurodevelopmental syndrome. Nature 2024 632:8026 632, (2024-07-11).

21. M. Irimia., A Highly Conserved Program of Neuronal Microexons Is Misregulated in Autistic Brains. Cell 159, (2014/12/18).

22. C. S. Leung., Dysregulation of the chromatin environment leads to differential alternative splicing as a mechanism of disease in a human model of autism spectrum disorder. Human Molecular Genetics 32, (2023/05/05).

23. S. Thacker., Alternative splicing landscape of the neural transcriptome in a cytoplasmic-predominant Pten expression murine model of autism-like Behavior. Translational Psychiatry 2020 10:1 10, (2020-11-06).

24. J. M. Lizcano., LKB1 is a master kinase that activates 13 kinases of the AMPK subfamily, including MARK/PAR-1. The EMBO Journal 23, (2004 Feb 19).

25. J. Courchet., Terminal Axon Branching Is Regulated by the LKB1-NUAK1 Kinase Pathway via Presynaptic Mitochondrial Capture. Cell 153, (2013/06/20).

26. V. Courchet., Haploinsufficiency of autism spectrum disorder candidate gene NUAK1 impairs cortical development and behavior in mice. Nature Communications 2018 9:1 9, (2018-10-16).

27. M. Lanfranchi., The AMPK-related kinase NUAK1 controls cortical axons branching by locally modulating mitochondrial metabolic functions. Nature Communications 2024 15:1 15, (2024-03-21).

28. G. Cossa., Localized Inhibition of Protein Phosphatase 1 by NUAK1 Promotes Spliceosome Activity and Reveals a MYC-Sensitive Feedback Control of Transcription. Molecular Cell 77, (2020/03/19).

29. i. A. Ilik., SON and SRRM2 are essential for nuclear speckle formation. eLife 9, (2020-10-23).

30. X. Zhu., Whole-exome sequencing in undiagnosed genetic diseases: interpreting 119 trios. Genetics in Medicine 17, (2015/10/01).

31. T. Takenouchi, K. Miura, T. Uehara, S. Mizuno, K. Kosaki, Establishing SON in 21q22.11 as a cause a new syndromic form of intellectual disability: Possible contribution to Braddock– Carey syndrome phenotype. American Journal of Medical Genetics Part A 170, (2016/10/01).

32. Mari J. Tokita., De Novo Truncating Variants in SON Cause Intellectual Disability, Congenital Malformations, and Failure to Thrive. The American Journal of Human Genetics 99, (2016/09/01).

33. G. Meyer-Dilhet, J. Courchet, In Utero Cortical Electroporation of Plasmids in the Mouse Embryo. STAR Protocols 1, (2020/06/19).

34. R. Schmieder, R. Edwards, Quality control and preprocessing of metagenomic datasets. Bioinformatics 27, (2011/03/15).

35. M. Martin, Cutadapt removes adapter sequences from high-throughput sequencing reads. EMBnet.journal 17, (2011/05/02).

36. A. Dobin., STAR: ultrafast universal RNA-seq aligner. Bioinformatics 29, (2013/01/01).

37. P. Danecek., Twelve years of SAMtools and BCFtools. GigaScience 10, (2021/01/29).

38. S. Anders, P. T. Pyl, W. Huber, HTSeq— a Python framework to work with high-throughput sequencing data. Bioinformatics 31, (2015/01/15).

39. M. I. Love., Moderated estimation of fold change and dispersion for RNA-seq data with DESeq2. Genome Biology 2014 15:12 15, (2014-12-05).

40. S. Shen., rMATS: Robust and flexible detection of differential alternative splicing from replicate RNA-Seq data. Proceedings of the National Academy of Sciences 111, (2014-12-23).

41. L. Kolberg., g:Profiler—interoperable web service for functional enrichment analysis and gene identifier mapping (2023 update). Nucleic Acids Research 51, (2023/07/05).

42. A. C. Bishop., A chemical switch for inhibitor-sensitive alleles of any protein kinase. Nature 2000 407:6802 407, (2000/09).

43. C. Zhang et., A second-site suppressor strategy for chemical genetic analysis of diverse protein kinases. Nature Methods 2005 2:6 2, (2005-05-20).

44. M. S. Lopez, J. I. Kliegman, K. M. Shokat, The Logic and Design of Analog-Sensitive Kinases and Their Small Molecule Inhibitors. Methods in Enzymology 548, (2014/01/01).

45. A. Zagórska., New Roles for the LKB1-NUAK Pathway in Controlling Myosin Phosphatase Complexes and Cell Adhesion. Science Signaling 3, (2010-03-30).

46. S. Banerjee., Characterization of WZ4003 and HTH-01-015 as selective inhibitors of the LKB1-tumour-suppressor-activated NUAK kinases. Biochemical Journal 457, (2014/01/01).

47. E.-Y. Ahn., SON Controls Cell-Cycle Progression by Coordinated Regulation of RNA Splicing. Molecular Cell 42, (2011/04/22).

48. J. Tapial., An atlas of alternative splicing profiles and functional associations reveals new regulatory programs and genes that simultaneously express multiple major isoforms. Genome Research 27, (2017-10-01).

49. S. M. Weyn-Vanhentenryck., Precise temporal regulation of alternative splicing during neural development. Nature Communications 2018 9:1 9, (2018-06-06).

50. J. E. O’Brien, M. H. Meisler, Frontiers | Sodium channel SCN8A (Nav1.6): properties and de novo mutations in epileptic encephalopathy and intellectual disability. Frontiers in Genetics 4, (2013/10/28).

51. J. E. O’Brien., Rbfox proteins regulate alternative splicing of neuronal sodium channel SCN8A. Molecular and Cellular Neuroscience 49, (2012/02/01).

52. N. W. Plummer, M. W. McBurney, M. H. Meisler, Alternative Splicing of the Sodium Channel SCN8A Predicts a Truncated Two-domain Protein in Fetal Brain and Non-neuronal Cells *. Journal of Biological Chemistry 272, (1997/09/19).

53. M. Ueda., Knockdown of Son, a mouse homologue of the ZTTK syndrome gene, causes neuronal migration defects and dendritic spine abnormalities. Molecular Brain 2020 13:1 13, (2020-05-24).

54. I. Iossifov., De Novo Gene Disruptions in Children on the Autistic Spectrum. Neuron 74, (2012/04/26).

55. I. Iossifov., The contribution of de novo coding mutations to autism spectrum disorder. Nature 2014 515:7526 515, (2014-10-29).

56. S. Alemany., New suggestive genetic loci and biological pathways for attention function in adult attention-deficit/hyperactivity disorder. American Journal of Medical Genetics Part B: Neuropsychiatric Genetics 168, (2015/09/01).

57. M. R. Johnson., Systems genetics identifies a convergent gene network for cognition and neurodevelopmental disease. Nature Neuroscience 2015 19:2 19, (2015-12-21).

58. D. Vojinovic., Genome-wide association study of 23,500 individuals identifies 7 loci associated with brain ventricular volume. Nature Communications 2018 9:1 9, (2018-09-26).

59. H. Cui., Arg-tRNA synthetase links inflammatory metabolism to RNA splicing and nuclear trafficking via SRRM2. Nature Cell Biology 2023 25:4 25, (2023-04-14).

60. K. A. Alexander., Nuclear speckles regulate functional programs in cancer. Nature Cell Biology 2025 27:2 27, (2025-01-02).

61. E. Matsumoto., AMP-activated protein kinase regulates alternative pre-mRNA splicing by phosphorylation of SRSF1. Biochemical Journal 477, (2020/06/26).

62. J. D. Kohtz., Protein–protein interactions and 5’-splice-site recognition in mammalian mRNA precursors. Nature 1994 368:6467 368, (1994/03).

63. J. Y. Wu, T. Maniatis, Specific interactions between proteins implicated in splice site selection and regulated alternative splicing. Cell 75, (1993/12/17).

64. J. F. Cáceres, T. Misteli, G. R. Screaton, D. L. Spector, A. R. Krainer, Role of the Modular Domains of SR Proteins in Subnuclear Localization and Alternative Splicing Specificity. Journal of Cell Biology 138, (1997/07/28).

65. N. Kataoka, J. L. Bachorik, G. Dreyfuss, Transportin-SR, a Nuclear Import Receptor for SR Proteins. Journal of Cell Biology 145, (1999/06/14).

66. C.-T. Sun., Transcription Repression of Human Hepatitis B Virus Genes by Negative Regulatory Element-binding Protein/SON *. Journal of Biological Chemistry 276, (2001/06/29).

67. J.-H. Kim., SON drives oncogenic RNA splicing in glioblastoma by regulating PTBP1/PTBP2 switching and RBFOX2 activity. Nature Communications 2021 12:1 12, (2021-09-21).

68. S. Nakahata, S. Kawamoto, Tissue-dependent isoforms of mammalian Fox-1 homologs are associated with tissue-specific splicing activities. Nucleic Acids Research 33, (2005/04/01).

69. G. S. Huh, R. O. Hynes, Regulation of alternative pre-mRNA splicing by a novel repeated hexanucleotide element. Genes & Development 8, (1994-07-01).

70. K. K. Kim, R. S. Adelstein, S. Kawamoto, Identification of Neuronal Nuclei (NeuN) as Fox-3, a New Member of the Fox-1 Gene Family of Splicing Factors *. Journal of Biological Chemistry 284, (2009/11/06).

71. A. J. M. Dingemans., Establishing the phenotypic spectrum of ZTTK syndrome by analysis of 52 individuals with variants in SON. European Journal of Human Genetics 2021 30:3 30, (2021-09-15).

