## Supplementary figures and methods for "The AMPK-related kinase NUAK1 regulates neuronal morphogenesis through the RNA splicing co-factor SON"

### SUPPLEMENTARY MATERIAL

This document contains:

- Supplementary figures 1 to 5
- Supplementary materials and methods
- References

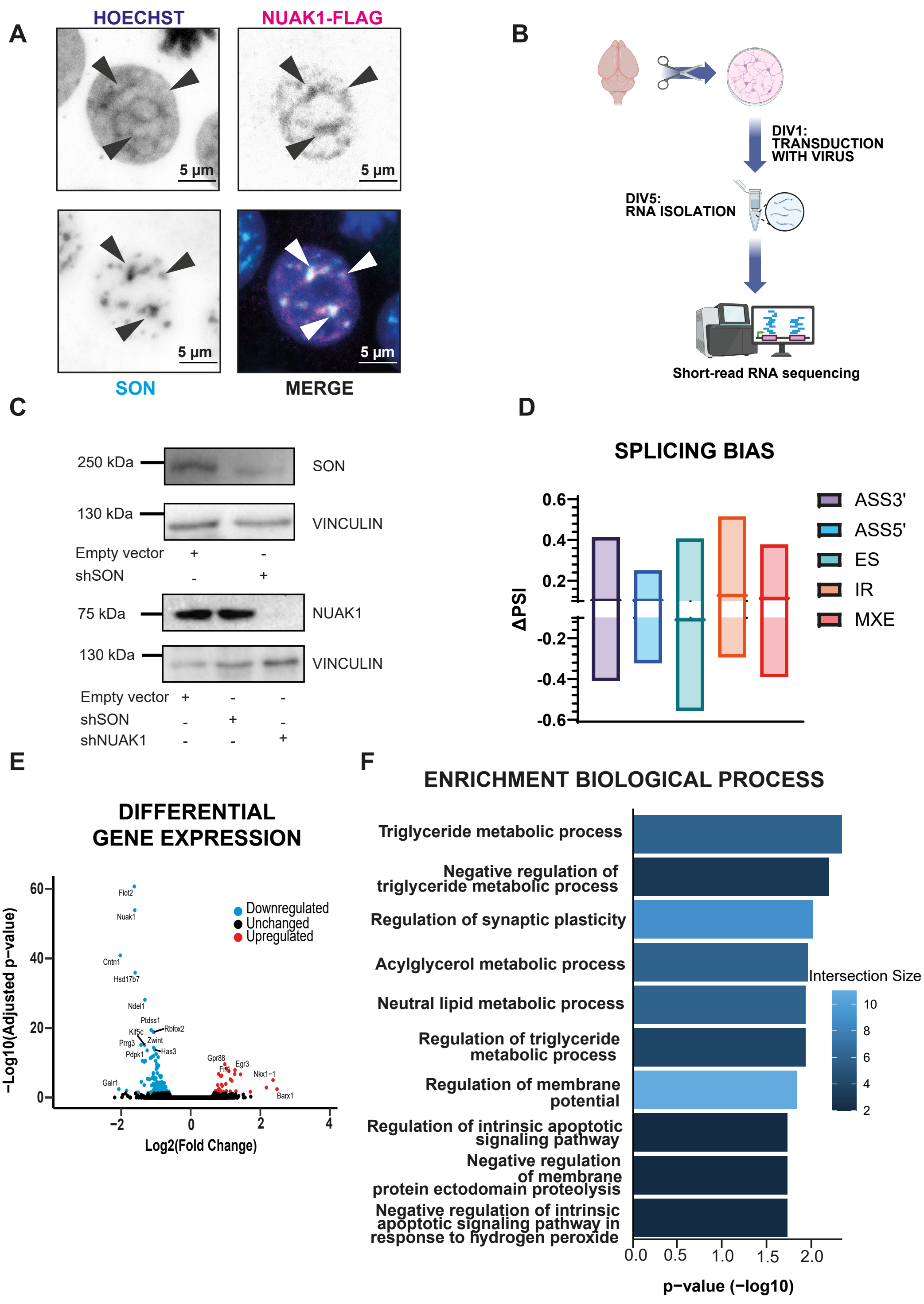


**Supplemental Figure 1.** (A) Immuno-fluorescence assay in 293T cells showing co-localization of Flag-tagged NUAK1 and nuclear speckle marker protein SON. (B) Schematic representation of experimental plan. (C) Validation of shRNA against *Nuak1* and *Son* in primary cortical neurons. (D) Splicing bias in splicing events in NUAK1-deficient neurons. The thick line indicates the median. (E) Volcano plot showing differential gene expression analysis (DGE) in NUAK1-deficient cortical neurons. Genes that were downregulated are represented in blue, while upregulated genes are shown in red. (F) gProfiler enrichment analysis for biological process on genes that were significantly differentially expressed in NUAK1-deficient cortical neurons.

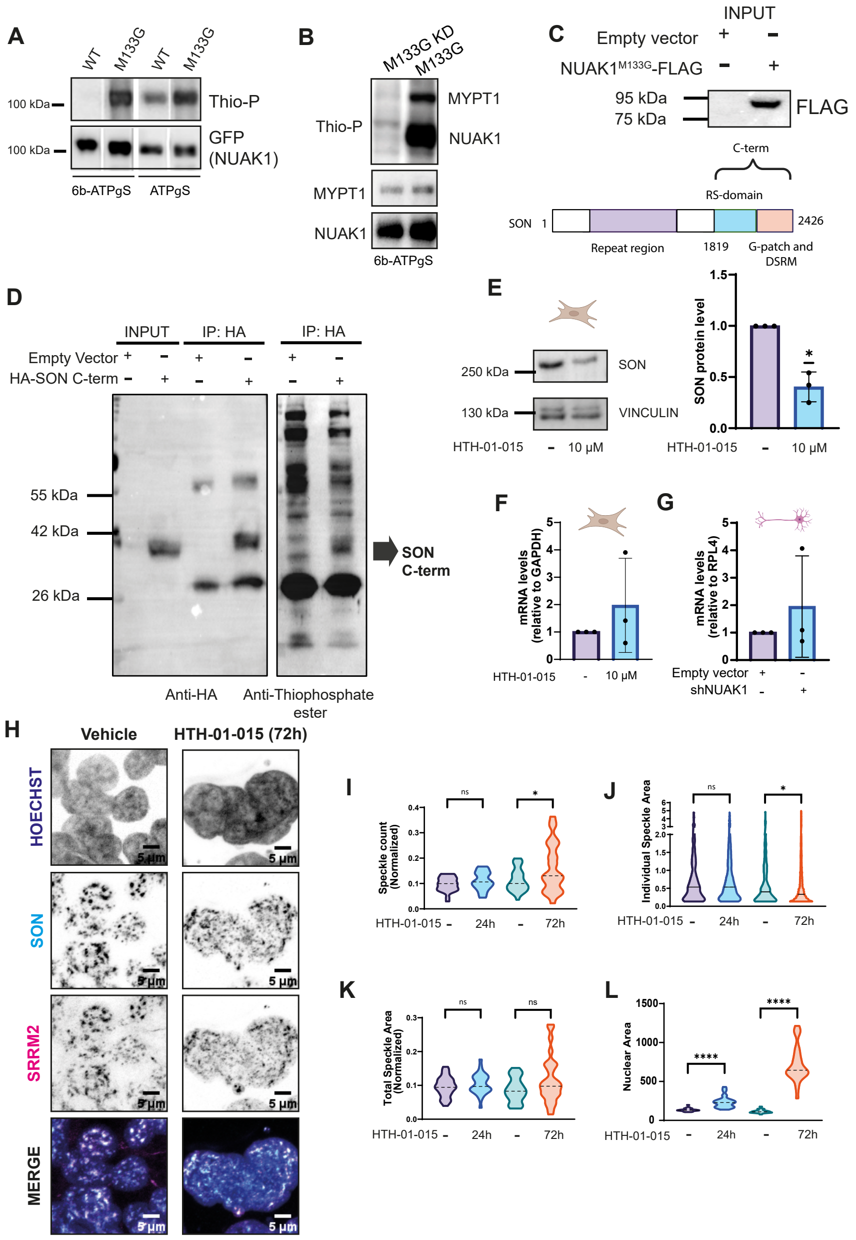


**Supplemental Figure 2.** (A) Western blot showing validation of the NUAK1$^{M133G}$ kinase assay. (B) Western blot validating that NUAK1$^{M133G}$ is able to phosphorylate a known NUAK1 target, i.e. MYPT1. (C) Western blot showing expression of NUAK1$^{M133G}$ for the kinase assay (up) and SON linear structure (down). (D) Western blot representing the analog sensitive kinase assay using Flag-NUAK1$^{M133G}$ detecting the phosphorylation of the HA-SON C-terminus. (E) Western blot representing protein levels of SON in 293T cells in the presence or absence of HTH-01-015, Paired t-test, * $p < 0.05$. (F-G) qPCR analysis showing *SON* mRNA levels in 293T cells and neurons, respectively. *GAPDH* was used as a housekeeping gene for 293T cells, while *Rpl4* was used for the primary cortical neurons. (H) Immuno-fluorescent images showing SON and SRRM2 in HEK293 cells in the presence or absence of NUAK1 inhibitor HTH-01-015 (72h incubation). (I-K) Nuclear speckle parameters based on SRRM2 immuno-fluorescence in 293T cells: speckle count (I; normalized to nuclear area), individual speckle area (J), and total speckle area (K; normalized to nuclear area), Mann-Whitney test, *** $p < 0.001$. (L) Nuclear area as measured by DAPI staining in HEK293 cells in the presence or absence of NUAK1 inhibitor HTH-01-015. Mann-Whitney test, **** $p < 0.0001$.

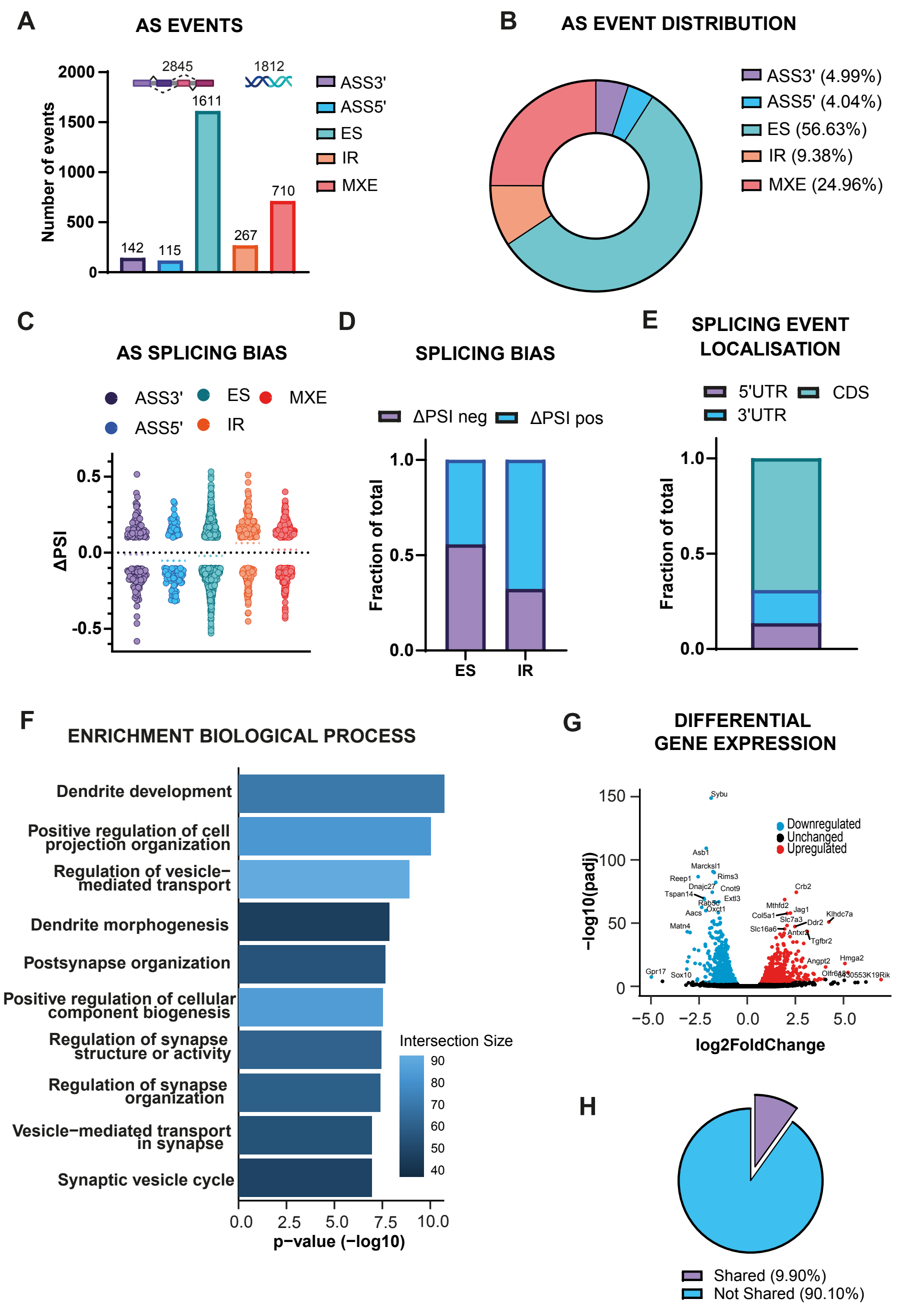


**Supplemental Figure 3.** (A) Number of alternative splicing (AS) events per category compared to the control condition in SON-deficient cortical neurons (5DIV): alternative splice site 3' (ASS3'), ASS5', exon skipping or inclusion (ES), intron retention (IR) and mutually exclusive exons (MXE). (B) AS event distribution per category in SON-deficient cortical neurons. (C) Scatter plot showing the distribution of Δ percent spliced in (PSI) per splicing event category in SON-deficient cortical neurons. (D) Splicing bias in splicing events in SON-deficient neurons as measured by the ratio of positive and negative ΔPSI. (E) Splicing event topology in SON-deficient cortical neurons. (F) gProfiler enrichment analysis for biological process on genes that were significantly differentially spliced in NUAK1-deficient cortical neurons. (G) Volcano plot showing differential gene expression analysis (DGE) in SON-deficient cortical neurons. Genes that were downregulated are represented in blue, while upregulated genes are shown in red. (H) Overlap between the genes found in the AS analysis and the genes found in the DGE analysis in SON-deficient cortical neurons.

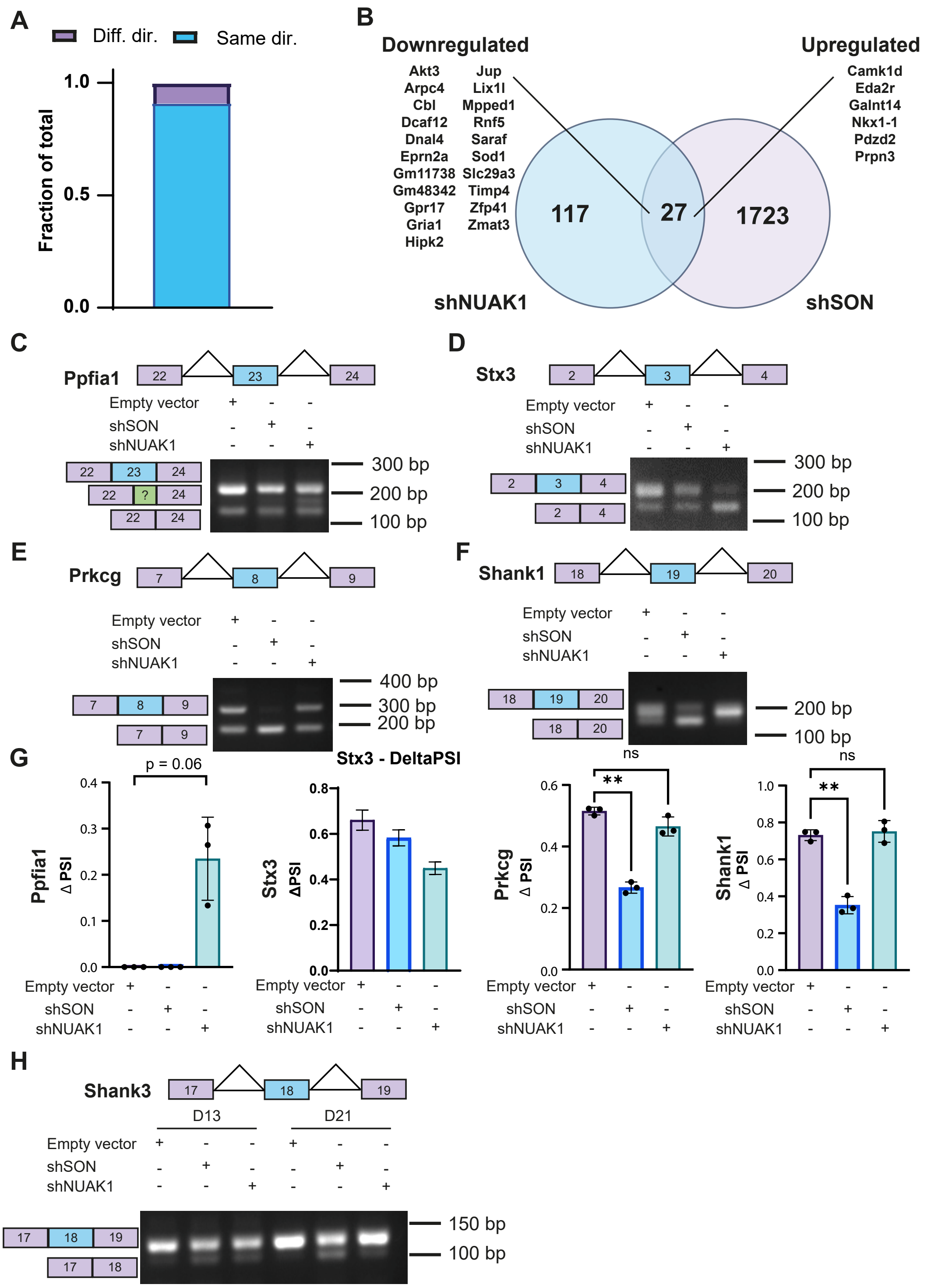


**Supplemental Figure 4.** (A) Comparison between ΔPSI of splicing events in the shared splicing signature between NUAK1- and SON-deficient cortical neurons going in the same or opposite direction. (B) Overlap between the DGE analysis in NUAK1- and SON-deficient cortical neurons. (C-D-G) Analysis and quantification of splicing events that are affected by NUAK1-deficiency in primary cortical neurons (*Ppfia1* and *Stx3*), One-way ANOVA. (E-F-G) Analysis and quantification of splicing events that are affected by SON-deficiency in primary cortical neurons (*Prkcg* and *Shank3*), One-way ANOVA. ** $p < 0.01$. (H) Developmental splicing pattern across 5DIV, 13DIV and 21 DIV of exon 18 in *Shank3*.

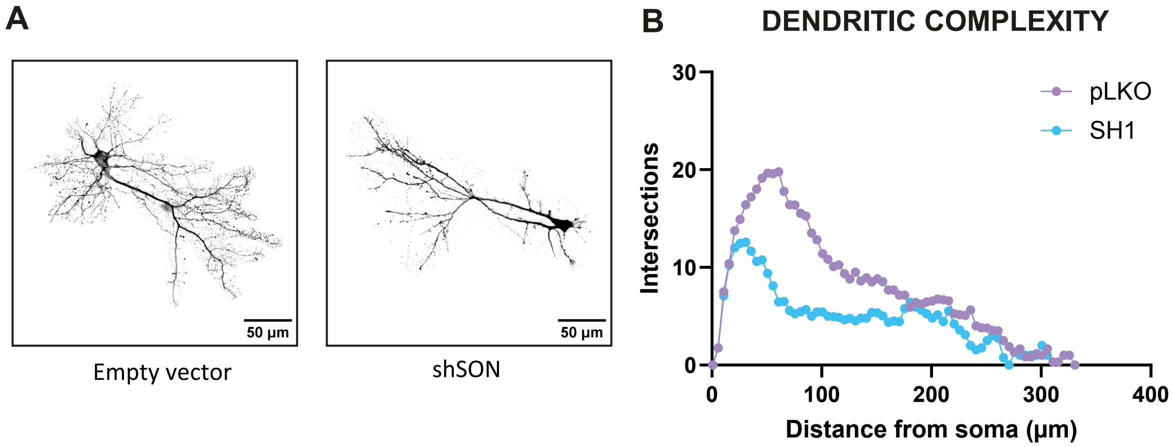


**Supplemental Figure 5.** (A) Representative images of primary cortical neurons at 13DIV. (B) Dendritic complexity as analysed by Sholl analysis in control cortical neurons *versus* SON-deficient cortical neurons (13DIV).

### SUPPLEMENTARY MATERIALS AND METHODS

*Plasmids*

To express fluorescent proteins, we used the following, previously described plasmids: mVENUS expressing vector pSCV2 (*1*), pCAG-mScarlet-I (*2*). Mouse NUAK1 coding sequence was cloned in a eGFP-C1 vector between the XhoI and EcoRI sites as described previously (*3*). A Flag-tag was added to the N-terminal extremity of the sequence by PCR. From this vector, we generated mutants M133G and K85A/M133G by site-directed mutagenesis using the Quikchange II site-directed mutagenesis kit (Stratagene) using the following primers (forward): 5'-CAAAGATAAGATTGTGATCATCGGGGAATATGCCAGCAAAGGAGAG-3' (M133G) and 5'-GCCGAGTGGTTGCTATAGCATCCATCCGTAAGGAC-3' (K85A). Flag-NUAK1 mutants were subsequently inserted into a pCIG2 (pCAG-IRES-eGFP version 2) vector between XhoI and EcoRI sites. pCIG2 vector was a kind gift from Dr. Franck Polleux (Columbia University, New York, USA).

The HA-tagged human SON C-term plasmid were described previously (*4*).

Empty vector pLKO.1 and shRNA targeting plasmids toward mouse NUAK1 (TRCN0000024112) from The RNAi Consortium shRNA Library (TRC) from the Broad Institute were described previously (*5*). shRNA targeting plasmids toward mouse SON (TRCN0000102292) was purchased from Sigma-Aldrich. All plasmids used in this study can be found in Table 1.

| ***Plasmid name*** | ***Backbone*** | ***Insert*** |
|---|---|---|
| *shSON (TRCN0000102292)* | *pLKO.1* | *CGCTCTATGATGTCTTATGAA* |
| *shNUAK1 (TRCN0000024112)* | *pLKO.1* | *GCAGAGAGAATCTGGCTACTA* |
| *pSCV2* | *pSilencer2.1* | *CAG-Venus-pA* |
| *ScarletRed* | *pCAG2* | *mScarlet-i* |
| *pCIG2-Flag-NUAK1* | *pCAG-IRES-GFP* | *mouse Flag-NUAK1* |
| *pCIG2-Flag-NUAK1$^{M133G}$* | *pCAG-IRES-GFP* | *mouse Flag-NUAK1$^{M133G}$* |
| *HA-SON C-terminus* | *pcDNA3.1* | *Human HA-SON isoform f siRNA-resistant 1816-2426* |
| *VSV G* | *pMD2.G* | *VSV G* |
| *psPAX2* | *psPAX2* | */* |

*Table 1: Plasmids*

*Cell lysis and Western blot*

Cortical neurons were lysed with sodium deoxycholate lysis buffer (50 mM Tris-HCl, 600 mM NaCl, 1% NP40, 0.5% sodium deoxycholate, protease inhibitor without EDTA (Roche, 11836170001), phosphatase inhibitor (Roche, 04906837001)) for 30 minutes on ice. Afterwards, cells were subjected to sonication (15 seconds on; 30 seconds off; 5 cycles) and centrifuged at 12,000 *g* for 10 min at 4 °C. Supernatant was collected and protein concentration was determined using DC Protein Assay (Biorad, Cat. #5000114). HEK293 cells were subjected to the same protocol except that they were lysed using NP40 lysis buffer (20 mM Tris-HCl pH 8.0, 150 mM NaCl, 1% NP40, 10% glycerol, protease inhibitor cocktail without EDTA (Roche, 11836170001) and phosphatase inhibitor (Roche, 04906837001)). Samples containing 30 µg of protein were loaded on a homemade polyacrylamide gel and for migration a Tris-Glycine-SDS buffer (Euromedex, EU0510) was used. Transfer (18h, 40V, 4°C) was done in a Tris-Glycine buffer (Euromedex, EU0550) on polyvinylidene fluoride membranes (0.45 µm). Membranes were subsequently blocked during 1h using a TBS-Tween 0.1% solution, supplemented with 5% bovine serum albumin (blocking buffer) and incubated overnight with primary antibody diluted in blocking buffer at 4°C (see Table 2). HRP-conjugated secondary antibodies (see Table 3) were diluted in blocking buffer as well and incubated for 1h in blocking before, followed by three washes (10 min each) with TBS-Tween 0.1%. Imaging of the membrane was done using ECL$^{TM}$ Prime Western Blotting Detection Reagent (Cytiva, RPN2236) according to the manufacturer's instructions using a Biorad Chemidoc$^{TM}$ MP Imaging System.

| ***Antibody*** | ***Reference*** | ***Company*** | ***Species*** | ***Dilution*** | ***Application*** |
|---|---|---|---|---|---|
| *Anti-SON* | *PA5-115947* | *Invitrogen* | *Rabbit* | *1:1000* | *Western blot* |
| *Anti-SON* | *HPA023535* | *Merck* | *Rabbit* | *1:1000* | *Immuno-fluorescence* |
| *Anti-Vinculin* | *V9131* | *Merck* | *Mouse* | *1:1000* | *Western blot* |
| *Anti-NUAK1* | *47677S* | *CST* | *Rabbit* | *1:1000* | *Western blot* |
| *Anti-SRRM2* | *S4045* | *Merck* | *Mouse* | *1:500* | *Immuno-fluorescence* |
| *Anti-GFP* | *600-901-B12* | *Rockland* | *Chicken* | *1:1000* | *Immuno-fluorescence* |
| *Anti-Flag (M2)* | *F3165* | *Merck* | *Mouse* | *1:1000* | *Western blot* |
| *Anti-Flag (M2)* | *F3165* | *Merck* | *Mouse* | *1:200* | *Immuno-fluorescence* |

| | | | | | |
|---|---|---|---|---|---|
| *Anti-Flag (M2)* | *F3165* | *Merck* | *Mouse* | *1:100* | *Immuno-precipitation* |
| *Anti-HA* | *901514* | *Biolegend* | *Mouse* | *1:100* | *Immuno-precipitation* |
| *Anti-HA* | *901514* | *Biolegend* | *Mouse* | *1:1000* | *Western blot* |
| *Anti-thiophos-phate ester* | *SD2020* | *Thermo-Fisher* | *Rabbit* | *1:1000* | *Western blot* |

*Table 2: Primary Antibodies*

| ***Antibody*** | ***Reference*** | ***Company*** | ***Dilution*** | ***Application*** |
|---|---|---|---|---|
| *Anti-rabbit IgG HRP-linked* | *7074S* | *CST* | *1:5000* | *Western blot* |
| *Anti-mouse IgG HRP-linked* | *7076S* | *CST* | *1:5000* | *Western blot* |
| *Alexa Fluor™ Goat anti-chicken IgY* | *A11029* | *Invitrogen* | *1:2000* | *Immuno-fluorescence* |
| *Alexa Fluor™ 488 Donkey anti-rabbit IgG* | *A21206* | *Invitrogen* | *1:2000* | *Immuno-fluorescence* |
| *Alexa Fluor™ 546 Donkey anti-rabbit IgG* | *A10040* | *Invitrogen* | *1:2000* | *Immunofluorescence* |
| *Alexa Fluor™ 647 Donkey anti-mouse IgG* | *A31571* | *Invitrogen* | *1:2000* | *Immunofluorescence* |

*Table 3: Secondary Antibodies*

*ATP analogue labelling of NUAK1 substrates*

To enable use of ATP analogues for direct labelling of NUAK1 substrates, we first identified the gatekeeper methionine of the ATP-binding pocket based on sequence homology to AMPK$\alpha$2 (*6*). Methionine 133 of HA-tagged murine Nuak1 was mutated to Glycine using a QuickChange II XL mutagenesis kit (Agilent), and subcloned into pBabe-Bleo expression vector. Stable cell lines were generated by infecting U2OS cells with pBabe-Nuak1$^{M133G}$ or pBabe-Nuak1$^{WT}$ retrovirus, followed by selection on Bleomycin. U2OS-Nuak1$^{M133G}$ and U2OS-Nuak1$^{WT}$ cells were cultured for 3 passages in SILAC DMEM (without arginine and lysine, Life Technologies) supplemented with 84 mg/l $^{12}C_6{}^{14}N_4$ L-arginine and 146 mg/l $^{12}C_6{}^{14}N_2$ L-lysine

(that we refer to as 'light', Sigma-Aldrich), or 84 mg/l $^{13}C_6{}^{15}N_4$ L-arginine and 175 mg/l $^{13}C_6{}^{15}N_2$ L-lysine (that we refer to as 'heavy', Cambridge Isotope Laboratories), 2% FBS and 8% 10 kDa dialyzed FBS (PAA). SILAC-labelled cells in 150 mm dishes were washed with 10 ml of room temperature PBS and aspirated. Kinase reactions were starting by adding 950 ul of warm (37°C) kinase buffer (20 mM HEPES, 100 mM KOAc, 5 mM NaOAc, 2 mM MgOAc, 1 mM EGTA, 10 mM $MgCl_2$, 30 µg/ml Digitonin, 0.1 mM PHET ATP (Biolog, P 026), 0.1 mM ATP, 5 mM GTP, 0.45 mM AMP, protease/phosphatase Complete inhibitors (Roche)), and plates were slowly incubated on a shaker for 20 min at room temperature. The kinase reaction was stopped by adding 40 ul 0.5 M EDTA to a final concentration of 20 mM. Cells were scraped and collected in tubes and lysates were sonicated at amplitude 4 for 30 sec each and centrifuged at 14,000 rpm for 10 min. Supernatants were transferred to new tubes and alkylating agent p-nitrobenzyl mesylate (PNBM, Abcam, ab138910) was added at a final concentration of 2.5 mM. Samples were incubated at room temperature for 30 min with shaking/rotation. To remove PNBM samples were run over PD-10 columns (Amersham) and eluted with TGN IP buffer (50 mM Tris pH 7.5, 200 mM NaCl, 1 % Tween, 0.2 % NP-40, phosphatase/protease inhibitors).
A protein assay was performed on the eluted fractions and equal amounts of lysates of equivalent NUAK1M133G and empty vector heavy and light samples were mixed (~8 ml total) and precleared with protein G beads for 1 h 4°C with rotation. Beads were pelleted for 5 min at 1000 rpm and precleared lysate eluate collected. Anti-Thiophosphate ester antibody (Abcam, ab133473) was crosslinked to protein G beads (100 ul beads slurry with 50 ug antibody) using dimethyl pimelimidate (DMP, Sigma D-8388) according to Abcam's protocol book (Chapter 8.6 "Procedure for cross-linking the antibody to the beads"). The protein G beads with crosslinked antibodies were washed and resuspended in TGN IP buffer and equally divided over the precleared lysates and incubated overnight at 4°C with rotation. The beads were washed with TGN IP buffer and interacting proteins eluted off the beads using low pH glycine then neutralised by addition of 1 M Tris pH 8.9 making the end concentration in the samples 0.1 M glycine, 20 mM Tris-HCL ~pH 8.

*Proteomics*

Differentially labelled samples were mixed in equal amounts using a label-swap replication strategy: NUAK1$^{M133G}$ SILAC-Heavy and empty vector SILAC-Light generates forward replicate; NUAK1$^{M133G}$ SILAC-Light and empty vector SILAC-Heavy generates reverse replicate. Sample volume was reduced by SpeedVac to ~10 µL. Proteins were denatured in Laemmli buffer 4x and separated by SDS-PAGE on NuPAGE 10% bis-tris gel. The gel was stained with colloidal Coomassie dye.
The region containing the bands of interest were excised from gel and washed twice with 50 mM ammonium bicarbonate, and 50 mM ammonium bicarbonate with 50% acetonitrile.

Proteins in the gel bands were then reduced using dithiothreitol (10mM at 54°C for 30 minutes) and subsequently alkylated with iodoacetamide (55mM at room temperature for 45 minutes). Gel pieces were washed again with 50 mM ammonium bicarbonate; 50 mM ammonium bicarbonate with 50% acetonitrile; and finally dehydrated using acetonitrile before drying in a SpeedVac. Trypsin (5 µg/mL trypsin gold; Promega in 25 mM ammonium bicarbonate) was added and incubated for 12 h at 35 °C. Tryptic peptides were extracted from gel pieces with two washes of 50% (v/v) acetonitrile/water + 1% trifluoroacetic acid (TFA) and subsequently dried in a SpeedVac. Dried peptides were resuspended in 0.1% TFA and desalted using StageTip (*7*).

Peptides resulting from digestion were separated by nanoscale C18 reverse-phase liquid chromatography performed on an EASY-nLC II (Thermo Scientific) coupled to a Linear Trap Quadrupole - Orbitrap Velos (Thermo Scientific). Elution was carried out at a flow rate of 200 nL/min using a binary gradient, into a 20 cm fused silica emitter (New Objective) packed in-house with ReproSil-Pur C18-AQ, 1.9 µm resin (Dr Maisch GmbH), for a total run-time duration of 70 minutes. Packed emitter was kept at 35 °C by means of a column oven (Sonation) integrated into the nanoelectrospray ion source (Thermo Scientific). An Active Background Ion Reduction Device (ABIRD) was used to decrease air contaminants signal level.

The Orbitrap Velos was operated in positive mode using a spray voltage, 2.4 kV, and an ion transfer tube temperature of 200°C. The mass spectrometer was used in data-dependent acquisition mode (DDA). A full scan (FT-MS) was acquired at a target value of 1,000,000 ions with resolution R = 60,000 over mass range of 350-1600 amu. The top ten most intense ions were selected for fragmentation in the linear ion trap using Collision Induced Dissociation (CID) using a maximum injection time of 25 ms or a target value of 5000 ions. Multiply charged ions from two to five charges having intensity greater than 5000 counts were selected through a 2 amu window and fragmented using normalized collision energy of 36 for 10ms. Former target ions selected for MS/MS were dynamically excluded for 20s. MS data were acquired using the XCalibur software (Thermo Fisher Scientific).

The MS Raw data were processed with MaxQuant software (*8*) version 1.5.0.36 and searched with Andromeda search engine (*9*), querying UniProt (*10*). First and main searches were performed with precursor mass tolerances of 20 ppm and 4.5 ppm, respectively, and MS/MS tolerance of 20 ppm. The minimum peptide length was set to six amino acids and specificity for trypsin cleavage was required. Cysteine carbamidomethylation was set as fixed modification, whereas Methionine oxidation and N-terminal acetylation were specified as variable modifications. The peptide, protein, and site false discovery rate (FDR) was set to 1%. For SILAC quantitation in MaxQuant, multiplicity was set to 2 and Arg0/Arg10, Lys0/Lys8 were used for ratio measurement of SILAC labelled peptides.

MaxQuant proteinGroups.txt output was further processed and analysed using Perseus software version 1.6.15.0 (*11*). The common reverse and contaminant hits (as defined in MaxQuant output) were removed. Only protein groups identified with at least one uniquely assigned peptide were used for the analysis.

| ***Gene*** | ***VastDB event*** | ***Forward primer*** | ***Reverse primer*** |
|---|---|---|---|
| *Scn8a* | *MmuEX0041275* | *GACCATCCGTACCATCCTGGA* | *GGTTAACGCCCATGATGCTGA* |
| *Shank3* | *MmuEX0042256* | *GTCCAAGTCCATGACGGCTG* | *AGTCAGCATCTGCAATGTCCG* |
| *Plch1* | *MmuEX0035625* | *CTACCGGCATGTCTACCTGGA* | *TTTCTGAAGAAGCGTGCCTGG* |
| *Arhgef10* | *MmuEX0005943* | *GAACGGCGAAGAAGGTGGAAA* | *TGAGAGACCAAGGACCTGGTC* |
| *Cadm1* | *MmuEX0008870* | *CGACGACAGAACCAGCAGTT* | *GCCCAGAATGATGAGCAAGCA* |
| *Atp2b1* | *MmuEX7004934* | *CGTGGCCAGATCTTGTGGTTT* | *TTTGTTAGGAGAGGGCGGAGG* |
| *Fxr1* | *MmuEX0019782* | *ACGGAGGACTGATGAAGATGC* | *CCCAGAGTACGCGGTAGCTTA* |
| *Ppfia1* | *MmuEX0036506* | *CATCAGCAACCCCCTGCATAG* | *CCACTCATTGCCGATCCACTC* |
| *Stx3* | *MmuEX0045595* | *TCGGAGCATGTAGAGGAAGCT* | *CTTCAGCTTGTTCCGGACGTT* |
| *Shank1* | *MmuEX0042245* | *CATCTCCCTGCGTTCCAAGTC* | *GCCATCTGATACACAGTCCGC* |
| *Prkcg* | *MmuEX0037125* | *TGAAGCCAGGGGATGTAGAGC* | *TCATACAATTCCAAGGGGTAATTGCA* |

*Table 4: PCR primers*

| ***Gene*** | ***Forward primer*** | ***Reverse primer*** |
|---|---|---|
| *SON (human)* | *TGTGTGCTAAGGCTGGTGTC* | *GGAGGTGCAGGCTTTAGGTT* |
| *GAPDH (human)* | *CAGGAGAGTGTTTCCTCGTCC* | *TTTGCCGTGAGTGGAGTCAT* |

*Table 5: qPCR primers*

*Viral production and transduction*

293T cells were seeded 24h before transfection at a density of 750 000 cells in a 6 cm plate. Subsequently 293T cells were transfected with a mix of the plasmid of interest (Control: pLKO.1 cloning vector), a plasmid expressing the viral envelope (pMD2.G; Addgene #12259) and a packaging plasmid (psPAX2; Addgene #12260) in a ratio 3:1:2, respectively, using a calcium phosphate transfection protocol. 24h after transfection, a medium change was performed and 48h later medium was collected, filtered (0.45 µm) and virus was precipitated with PEG-it virus precipitation solution (LV810A-1, System Biosciences) according to the manufacturer's instructions. The virus pellet was resuspended in sterile PBS 1x in 10 percent of the original volume. Neuronal cultures were transduced at a multiplicity of infection (MOI) of 5 at DIV1, DIV7 or DIV14 for harvesting at DIV5, DIV14 or DIV21, respectively. After 24h, the medium was changed.

### REFERENCES


1. R. Hand, F. Polleux, R. Hand, F. Polleux, Neurogenin2 regulates the initial axon guidance of cortical pyramidal neurons projecting medially to the corpus callosum. *Neural Development 2011 6:1* **6**, (2011-08-24).
2. M. Lanfranchi *et al.*, The AMPK-related kinase NUAK1 controls cortical axons branching by locally modulating mitochondrial metabolic functions. *Nature Communications 2024 15:1* **15**, (2024-03-21).
3. V. Courchet *et al.*, Haploinsufficiency of autism spectrum disorder candidate gene NUAK1 impairs cortical development and behavior in mice. *Nature Communications 2018 9:1* **9**, (2018-10-16).
4. J.-H. Kim *et al.*, SON and Its Alternatively Spliced Isoforms Control MLL Complex-Mediated H3K4me3 and Transcription of Leukemia-Associated Genes. *Molecular Cell* **61**, (2016/03/17).
5. J. Courchet *et al.*, Terminal Axon Branching Is Regulated by the LKB1-NUAK1 Kinase Pathway via Presynaptic Mitochondrial Capture. *Cell* **153**, (2013/06/20).
6. M. R. Banko *et al.*, Chemical genetic screen for AMPKα2 substrates uncovers a network of proteins involved in mitosis. *Molecular cell* **44**, (2011 Dec 1).
7. J. Rappsilber *et al.*, Protocol for micro-purification, enrichment, pre-fractionation and storage of peptides for proteomics using StageTips. *Nature Protocols 2007 2:8* **2**, (2007-08-02).
8. J. Cox, M. Mann, J. Cox, M. Mann, MaxQuant enables high peptide identification rates, individualized p.p.b.-range mass accuracies and proteome-wide protein quantification. *Nature Biotechnology 2008 26:12* **26**, (2008-11-30).
9. J. Cox *et al.*, Andromeda: A Peptide Search Engine Integrated into the MaxQuant Environment. (February 22, 2011).
10. UniProt: a worldwide hub of protein knowledge - PubMed. *Nucleic acids research* **47**, (01/08/2019).
11. S. Tyanova *et al.*, The Perseus computational platform for comprehensive analysis of (prote)omics data. *Nature Methods 2016 13:9* **13**, (2016-06-27).
